# A *TranSNP* in the DDIT4 mRNA can impact its translation efficiency and modulate p53-dependent responses in cancer cells

**DOI:** 10.1101/2025.04.02.646512

**Authors:** Meriem Hadjer Hamadou, Laura Alunno, Tecla Venturelli, Samuel Valentini, Davide Dalfovo, Francesca Lorenzini, Alessia Mattivi, Vincenza Vigorito, Glenda Paola Grupelli, Alessandro Matte’, Pamela Gatto, Michael Pancher, Chiara Valentini, Veronica De Sanctis, Roberto Bertorelli, Virginie Marcel, Emilio Cusanelli, Stefano Freddi, Giovanni Bertalot, Sara Zaccara, Marina Mione, Luca L. Fava, Alessandro Romanel, Alberto Inga

**Affiliations:** Laboratory of Transcriptional Networks; Laboratory of Bioinformatics and Computational Genomics; Laboratory of Experimental Biology; Armenise-Harvard Laboratory of Cell Division; Cell Biology and Molecular Genetics Laboratory; Cell Analysis and Separation Core Facility; High-Throughput Screening Core Facility; Next Generation Sequencing (NGS) Core Facility; Department of Cellular, Computational, and Integrative Biology, CIBIO, University of Trento, via Sommarive 9, 38123, Trento, Italy; Ribosome, Translation and Cancer » team, Centre de Recherche en Cancérologie de Lyon (CRCL), INSERM U1052, CNRS UMR5286, Centre Léon Bérard, Université de Lyon, Université Claude Bernard Lyon 1, 69008, Lyon, France; Department of Oncology and Hematology-Oncology, Università degli Studi di Milano, Milan, Italy; IEO, European Institute of Oncology IRCCS, Milan, Italy; Unità Operativa Multizonale di Anatomia Patologica, APSS, Trento, Italy, Centre for Medical Sciences – CISMed, University of Trento, Italy; Department of Systems Biology, Columbia University Irving Medical Center, NY, USA

**Keywords:** Polysome-profiling, translation, DDIT4, mTOR, p53, ER-stress

## Abstract

Relatively few studies have examined the link between SNPs and mRNA translation, despite the established importance of translational regulation in shaping cell phenotypes. We developed a pipeline analyzing the allelic imbalance in total and polysome-bound mRNAs from paired RNA-seq data of HCT116 cells and identified 40 candidate tranSNPs, i.e. SNPs associated with allele-specific translation. Among them, the SNP rs1053639 (T/A) on DNA damage-inducible transcript 4 (DDIT4) 3’UTR was identified, with the reference T allele showing a higher polysome association. rs1053639 TT clones generated by genome editing exhibited significantly higher DDIT4 protein levels than AA ones. The difference in DDIT4 proteins was even greater when cells were treated with Thapsigargin or Nutlin, two perturbations that induce DDIT4 transcription. The RNA-binding protein RBMX influenced these allele-dependent differences in DDIT4 protein expression, as shown by RNA-EMSA, RIP, and smiFISH assays. RBMX depletion reduced DDIT4 protein in TT clones to the AA levels. Functionally, TT clones more effectively repressed mTORC1 under ER stress, while AA clones outcompeted TT clones in vitro or when injected in zebrafish embryos. RBMX depletion increased the fitness of TT cells in co-culture experiments. The rs1053639 AA genotype, under a recessive model, correlates with poor prognosis in TCGA cancer data.

**Key points:**

- Translatome analysis in HCT116 cells revealed allele-specific mRNA translation for 40 SNPs
- rs1053639 (T/A) in DDIT4 3’UTR showed allelic differences in mRNA localization & protein expression
- AA cells showed weaker mTOR inhibition & higher proliferation; AA individuals had poorer prognosis

## Introduction

Many studies in the last decade have demonstrated the importance of translational regulation in shaping cell phenotype (1–4). Instead, relatively few studies focused on the identification and characterization of genetic variants impacting directly on mRNA translation, although allele-specific expression finely modulating epigenetic and transcriptional regulation has been revealed as critical for many pathophysiological processes (5–8). To investigate instances of allele-specific mRNA translation efficiency, we recently developed a method that focuses on single nucleotide polymorphisms (SNP) and integrates information from SNP genotyping with RNA-seq data performed on both total and polysome-bound mRNAs (9). Briefly, heterozygous SNPs mapped by genotyping assays are followed in RNA-seq data by calculating allelic fractions (AFs) for both total and polysomal RNA. Coverage threshold, consistency among biological replicates, and the differences in AF between paired total and polysomal RNA samples are used to nominate candidate tranSNPs. The method assumes that the translatome, defined as the ensemble of transcripts that co-sediment with polysomes on sucrose gradients, represents a proxy for the proteome (10). Also, it assumes that if two alleles expressed by a cell can be distinguished due to heterozygous SNPs or other types of variants, a difference in their relative abundance within the polysomes would signal an instance of allele-specific difference in translation efficiency. Hence, the name tranSNPs is attributed to the variants emerging from our pipeline (9). Besides polysomal profiling, Ribo-seq is another approach typically used to characterize the translatome. The method can reach a single-nucleotide positional resolution of the ribosomes along a transcript. The polysome-seq variant protocol could be potentially applied to our scope, as 80S monosomes, whose actual translational status is still somewhat debated (11), are removed before processing the samples by RNAse treatment to retrieve ribosome-protected fragments (12). However, we decided to use polysomal profiling to increase the possibility of obtaining robust coverage also in 5’- and 3’-UTR regions of mRNAs. Our earlier study identified 147 tranSNPs, corresponding to about 4% of analyzable SNPs in a dataset obtained in the MCF7 cancer cell lines. Of those, 39 were located in the 3’UTR (9) and could impact, directly or through structural features, the binding of trans-factors that, in turn, can affect translation efficiency or mRNA stability.

We have now extended our analysis using data produced in the HCT116 p53 wild-type cell line cultured in control conditions or treated with the MDM2 inhibitor Nutlin (13). While tranSNPs could potentially impact multiple cellular phenotypes and outcomes, we decided to continue to focus on a non-genotoxic p53-activating treatment in cancer cells due to profound changes in mRNA translation in cancer (4), and the growing evidence linking the tumor suppressor p53 to various features of mRNA translation control. For example, p53 was shown to modulate eIF4EBP1, mTORC1, and Endoplasmic Reticulum (ER) stress (14–16), and, through the modulation of Ribosomal Proteins (RBPs) and microRNAs, to impact the fate of specific sets of mRNAs (17–19). We identified 40 candidate tranSNPs, corresponding to about 7% of analyzable SNPs. Among these tranSNP candidates, four were prioritized for validation experiments considering minor allele frequency, the relevance of the associated genes in cancer, the position along the gene sequence, and the presence of predictions for overlapping trans factor binding sites. One candidate, rs1053639 T/A in the 3’UTR of the DDIT4 (REDD1, RTP801) gene, caught our particular interest. DNA damage-inducible transcript 4 DDIT4 is a negative modulator of mTORC1, is an established target of p53, RFX7, and ATF4, and acts as a tumor suppressor gene (20,21). We generated HCT116 cell line trios for rs1053639 exploiting a CRISPR/Cas9-mediated knock-in strategy (22) starting from the heterozygous (AT) parental cells. Homozygous TT and AA cell clones were characterized, which revealed differences in DDIT4 protein expression both in basal and induced conditions, alterations in the relative expression of mTOR markers, and changes in mRNA subcellular localization and translatability. Mechanistically, we implicated the RNA binding protein RBMX in a distinct fate of DDIT4 mRNA for rs1053639 alleles. We also established higher relative proliferation and fitness for A-homozygous cells in both *in vitro* and xenograft experiments in zebrafish embryos and explored the correlation between rs1053639 alleles and DDIT4 protein expression in primary tumor samples and the prognostic value of this SNP using TCGA data.

## Results

### TranSNPs can be identified in mock and Nutlin-treated HCT116 cells

Single-end RNA-seq data obtained from HCT116 cells under either mock or 10 µM Nutlin treatment for 16 hours were processed starting from row files by trimming, aligning, and deduplication steps (see Methods). SNPs in exonic and UTR regions were extracted from dbSNP. A list of heterozygous SNPs was compiled based on AF values between 0.2 and 0.8 in at least one of the four replicates in one of the two fractions sequenced (total and polysomal RNA), setting a minimal threshold of 20 reads for AF calculations (**Figure S1**). In addition, only the SNPs with AF measures available for all four biological replicates, the two fractions, and across the two treatments were considered to calculate changes in AF between the fractions (N= 582, **Table S1**). For each SNP, we exploited the concordance among differential AF measures across pairs of polysomal and total RNA biological replicates, performing paired statistics (**Figure S1**). This approach identified 40 putative tranSNPs (**Figure 1A**), of which 21 are located in 3’UTRs (**Table S2**). Overall, we extended the methodology previously developed (9) and observed that in HCT116, about 11% of genes with analyzable heterozygous SNPs show a difference in AF between paired total and polysome-bound mRNAs, suggesting allele-specific post-transcriptional and translational control.

**Figure 1.**
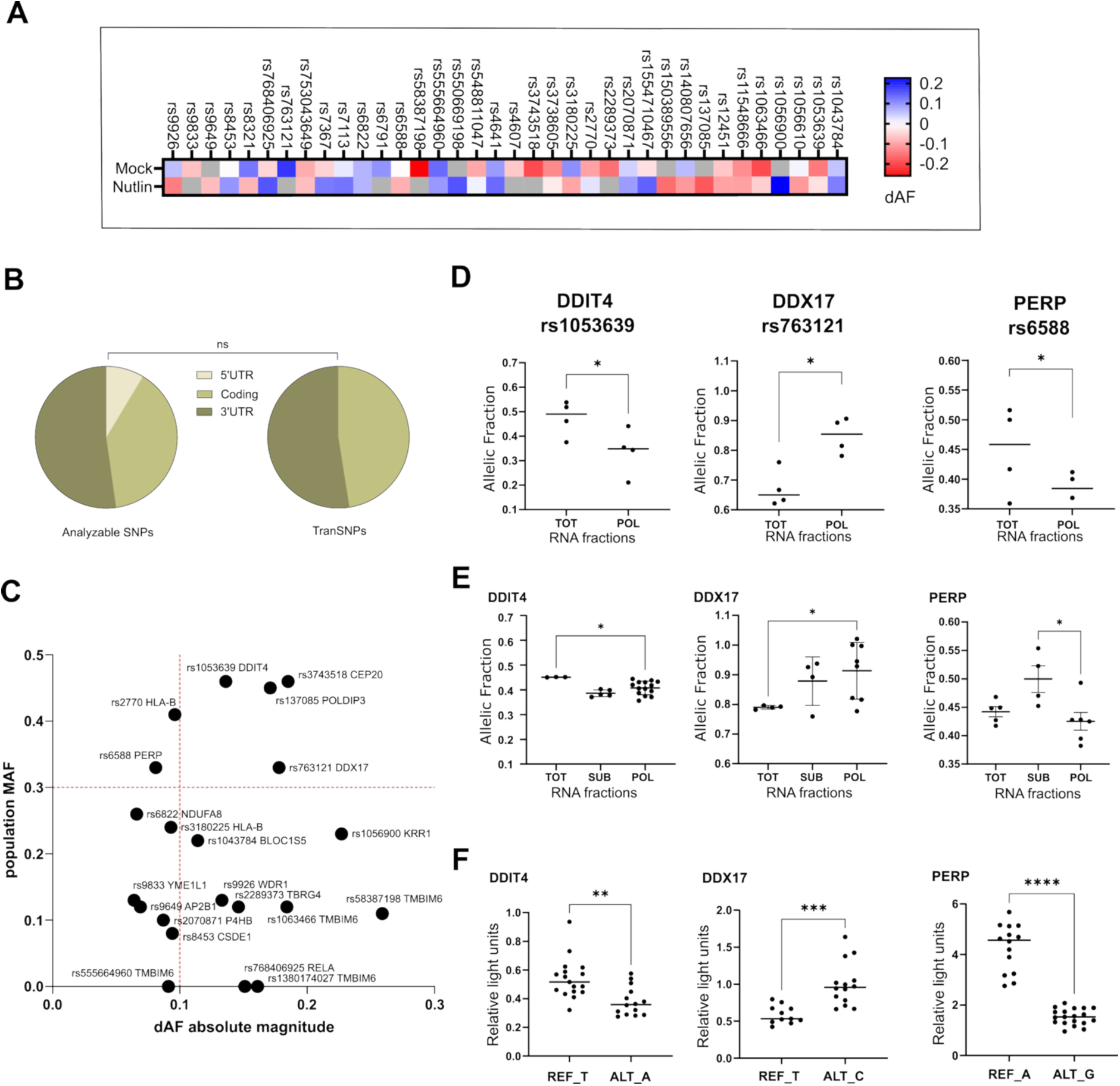
The tranSNPs in the DDIT4, DDX17, and PERP genes are validated by orthogonal approaches. **A**) Heatmap view of average differences in allelic fractions (AF) in the comparison between polysome-associated and total RNA for the 40 tranSNPs identified from the RNA-seq data from either mock or Nutlin-treatment conditions (Tables S1 and S2 present the AF value for each biological replicate; accession number for the RNA-seq data: GSE95024). AF was computed as the number of reads for the alternative allele over the sum of reads of the alternative and reference alleles. **B**) Distribution across coding and regulatory regions of the transcripts for all analyzable SNPs (Left) and for the identified tranSNPs (Right). The result of Chi-square contingency test is shown. **C**) Plot presenting the absolute delta Allelic Fraction (dAF) and the Minor Allele Frequency (MAF) for the identified tranSNPs located in 3’UTRs. MAF values were derived from the 1000 Genomes Project Phase 3, all individuals. The SNP rsID and associated gene names are shown. **D**) Plots of the AF for the rs1053639 (in DDIT4), rs763121 (in DDX17), and rs6588 (in PERP) tranSNPs obtained through RNA-seq of cytoplasmic total (TOT) and polysome-bound mRNAs (POL). Individual replicates and the AF mean are plotted. *p-value < 0.05, two-tailed, paired t-test. **E)** AF values of the selected tranSNPs calculated from Sanger sequencing electropherograms, comprising also the subpolysomal -RNP to 80S-(SUB) fractions. Individual replicates and the AF mean are plotted. *p-value < 0.05, two-tailed, paired t-test. **F)** Results of the dual luciferase assays presenting the relative activity (average and individual biological replicates) for the reference (REF_) and alternative (ALT_) alleles for each of the three selected tranSNPs cloned separately into the pGL4.13 vector immediately downstream of the Firefly luciferase stop codon. Data was normalized using Renilla luciferase. **p-value < 0.01. ***p-value < 0.001. ****p-Value < 0.0001, two-tailed, unpaired t-test.

Candidate tranSNPs for downstream analyses and validation were chosen considering several SNP features (**Figure 1B, 1C**, **Table S1**, **S2**) and information about the associated genes. Higher priority was given to SNPs with high magnitude of the AF imbalance, high minor allele frequency (MAF), and positioned in 3’UTR of genes whose expression is altered in cancer or whose function is associated with cancer-relevant features, especially p53-related. We selected a total of four SNPs for validation (**Figure 1D-F, S1**). First, we sought to confirm differential allele-specific expression found in polysomes versus total RNA. Sucrose gradient density sedimentation of cytoplasmic lysates was performed (10), and fractions were collected and analyzed by UV absorbance. RNA was recovered from the combined polysomal fractions and fractions corresponding to ribosomal subunits and 80S monosomes. Total RNA was also extracted. Amplicons were generated using primers specific for sequences surrounding the chosen SNPs (**Table S4**), Sanger sequenced, and the electropherograms were analyzed, expressing the results as AF (alternative allele signal over the sum of alternative + reference allele signals), as done for the RNA-seq data. For three out of the four chosen SNPs, an imbalance consistent in direction with what was measured in the RNA-seq dataset was found (**Figure 1E, S1D, S1F**).

We also used a targeted re-sequencing approach to measure allelic fractions from total and polysome-associated mRNAs, obtaining results consistent with the RNA-seq and Sanger sequencing (**Figure S1G**). Next, we asked whether the candidate tranSNPs could exhibit allele-specific differences in activity when extracted from their native context. To this aim, we cloned ∼500 nt fragments of the 3’UTR sequences harboring those SNPs downstream of a luciferase plasmid reporter, using HCT116 genomic DNA. Given the heterozygous state of the cell line, we retrieved clones for both alleles for each SNP. Dual luciferase assays were then performed in HCT116 cells, with results further supporting the observation that the SNPs and their immediate surroundings are sufficient to show allele-specific differences in reporter activity (**Figure 1F, S1**).

### Editing of HCT116 cells to obtain the three genotypes of rs1053639 tranSNP

It is worth noting that while the heterozygous state and the comparison of AF among two RNA compartments (total versus polysomal) are necessary to enable reliable identification of tranSNPs in our approach, we expect that the homozygous state for alternative and reference alleles would reveal the highest level of phenotypic impact of this category of genetic variants. Hence, our next step was constructing potentially isogenic cell clones by CRISPR/Cas9-mediated editing. We exploited the CRISPR/Cas9-mediated knock-in delivering the Cas9 and the guide RNA (gRNA) as a ribonucleoparticle (RNP) to reduce its off-target potential (22) (**Figure S2A**). Based on the collective RNA-seq, validation data, editing amenability, and the relevance of the gene in cancer, including its functional links with p53 (20,23,24), the T/A rs1053639 in DDIT4 was a promising tranSNP. HDR efficiency by the ICE Synthego analysis web tool (26) ranged from 5% to 9% for the initial bulk populations. 85 clones were isolated and characterized by Sanger sequencing. We obtained at least four clones homozygous for the rs1053639 T allele and four homozygous for the A allele. However, three clones (#3 TT, #5, and #6 AA) contained the same GGT 3 nt deletion, starting three nucleotides after the SNP site. We also obtained two clones bearing either a 9-nucleotide deletion (#7) or a 29-nucleotide deletion (#8) encompassing the SNP position (**Table S6**, **Figure S2B**, and methods). We considered at least two clones among the rs1053639 TT (#2, #3, #12, #13), and AA (#5, #6, #7, #10) along with the unedited AT clone (#1) to overcome the potential influence of clonal selection and adaptation and the above-mentioned editing limitations. We then used these cell models to study the SNP effect on various phenotypic assays.

### Higher DDIT4 protein levels in rs1053639 TT homozygous clones

First, we tested the impact of the tranSNP status on DDIT4 steady-state protein levels (**Figure 2A, B**). TT clones exhibited significantly higher relative protein expression than AA clones, with the AT cells intermediate. This result was consistent with the change in AF from RNA-seq or Sanger sequencing that showed lower A allele representation in the polysomal fraction. The difference in protein levels was not dependent on changes in protein half-life (**Figure 2C**), nor on mRNA levels and turnover (**Figure 2D**). An antisense DDIT4-AS1 transcript corresponding to the 3’UTR region was reported to affect the stability of the DDIT4 transcript (27,28). However, DDIT4-AS1 was equally expressed across TT and AA clones, and its expression was significantly lower than that of the DDIT4 transcript (**Figure 2E**). Next, we challenged the cells with Thapsigargin (100nM for 4 hours) (29,30), an ER stressor (31) that is a strong inducer of DDIT4 expression (**Figure 2F**). As expected, the treatment led to the marked induction of DDIT4 protein levels (**Figure 2G, 2H**). Notably, the difference in expression associated with the SNP genotype was maintained, and the lower expression of DDIT4 in AA clones was even more apparent. DDIT4 is also a recognized transcriptional target of p53, directly or through the transcription factor RFX7 (20,24). Hence, we examined whether the rs1053639 alleles differed in DDIT4 expression, also in response to p53 activation. Results showed that while the MDM2 inhibitor Nutlin led to comparable induction of p53 protein across the HCT116 DDIT4 clones, the associated induction in DDIT4 protein was higher for the TT homozygous clones and intermediate for the AT cells (**Figure S3A-S3C**). AA clones did not show significant DDIT4 induction in response to Nutlin.

**Figure 2.**
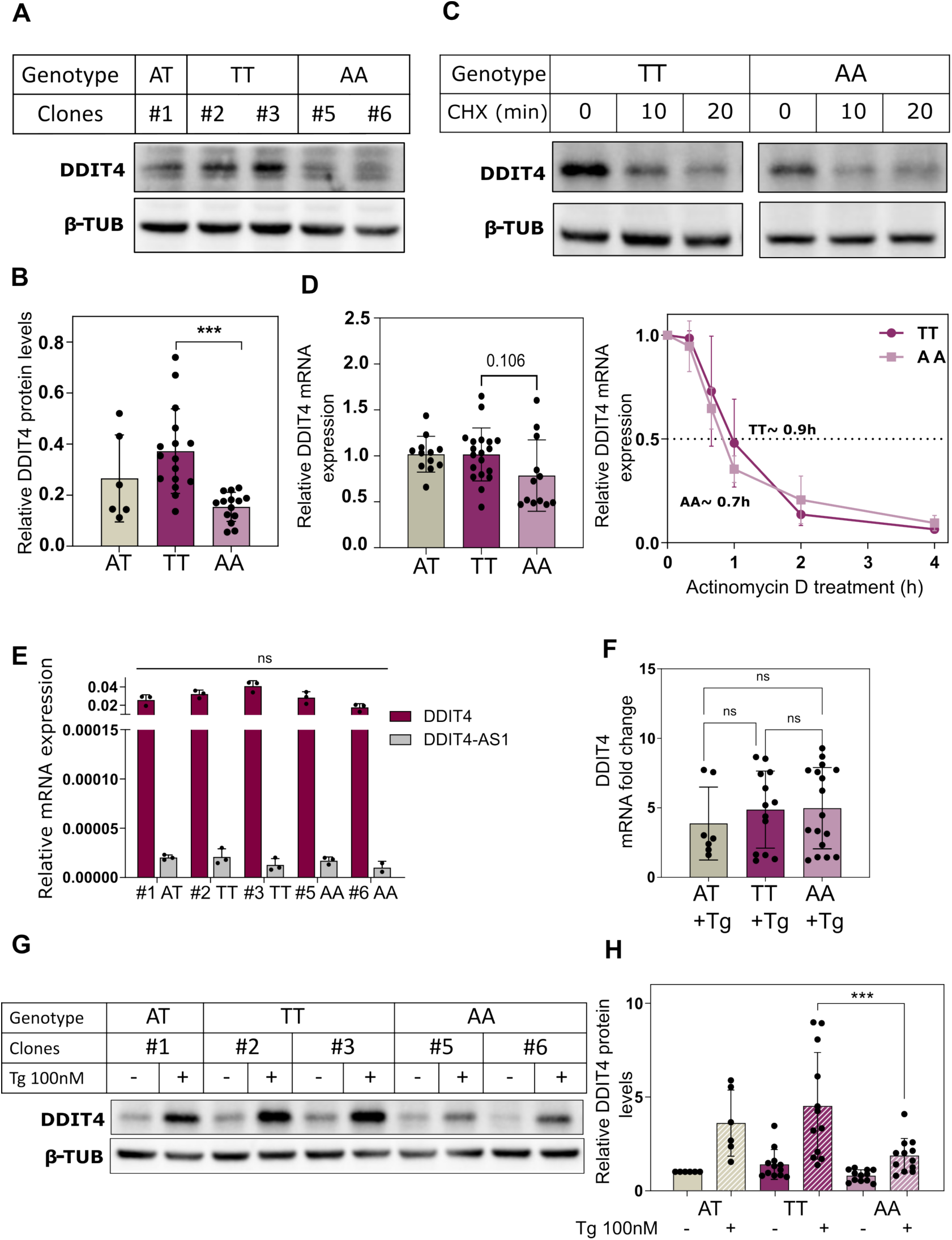
rs1053639 tranSNP impacts DDIT4 protein expression in HCT116-derived cells with the three genotypes. HCT116 cells were edited by CRISPR/Cas9 RNPs and donor DNA to obtain cell trios with the three genotypes for the rs1053639 tranSNP located in the proximal portion of the DDIT4 3’UTR. **A)** Western blot analysis of DDIT4 protein expression in HCT116 cells (A/T heterozygous) and DDIT4 rs1053639 homozygous clones (TT, AA). β-Tubulin was used as a reference. **B)** Densitometry analysis of the blots from several biological replicates shows a significant difference in relative DDIT4 protein abundance with the different tranSNPs genotypes. ***P-value <0.001, Ordinary one-way ANOVA. **C)** Western blot analysis of DDIT4 protein in HCT116 TT and AA homozygous clones for the rs1053639 treated with 100µg/ml cycloheximide (CHX) at the indicated time points. β-Tubulin was used as a reference. **D)** (Left) Relative DDIT4 mRNA expression as measured by RT-qPCR and normalized to GAPDH and YWHAZ. The Student’s t-test showed no significant differences in DDIT4 mRNA expression in the different clones. ns, not significant. (Right) Estimation of DDIT4 mRNA stability following inhibition of transcription by actinomycin D treatment (10µg/ml). The curve presents the residual relative DDIT4 mRNA levels (average and standard deviation of at least three biological replicates) at various treatment time points measured by RT-qPCR decay from HCT116 clones harboring TT or AA genotypes. The estimated half-lives of the DDIT4 mRNA in TT and AA homozygous clones are indicated. **E)** DDIT4-AS1 and DDIT4 mRNA levels measured by RT-qPCR and normalized to GAPDH across TT and AA clones. The expression of DDIT4-AS1 was negligible compared to the expression of the sense DDIT4 mRNA in our cellular models. Data represents the mean ± SD of three independent experiments. ns, not significant. **F)** HCT116 clones were exposed to 100 nM thapsigargin (Tg) or vehicle for 4 hours, and the induction of DDIT4 mRNA was measured by RT-qPCR with clones of the three rs1053639 genotypes. Average, standard deviation, and individual replicates are presented. ns, not significant, student’s t-test. **G-H**) Western blot and densitometry analysis showing relative DDIT4 protein induction upon thapsigargin (Tg 100nM) treatment of the three rs1053639 genotypes. β-Tubulin served as a control. ***p-value <0.001, unpaired t-test.

Since we observed higher polysomal association and higher DDIT4 protein levels from the rs1053639 T over the A allele, we examined at higher resolution the relative DDIT4 mRNA distribution across the sucrose gradient. For this experiment, RNA was recovered from ten individual fractions along the polysome profile (fractions 3 to 12, **Figure S4A**), and the DDIT4 mRNA distribution, along with CDKN1A and RPS26 controls, was measured by qPCR. Results were plotted as relative abundance, as previously described (32). In the mock condition, no significant differences in apparent translation efficiency for the DDIT4 mRNA between TT and AA homozygous cells were observed (**Figure 3A**).

**Figure 3.**
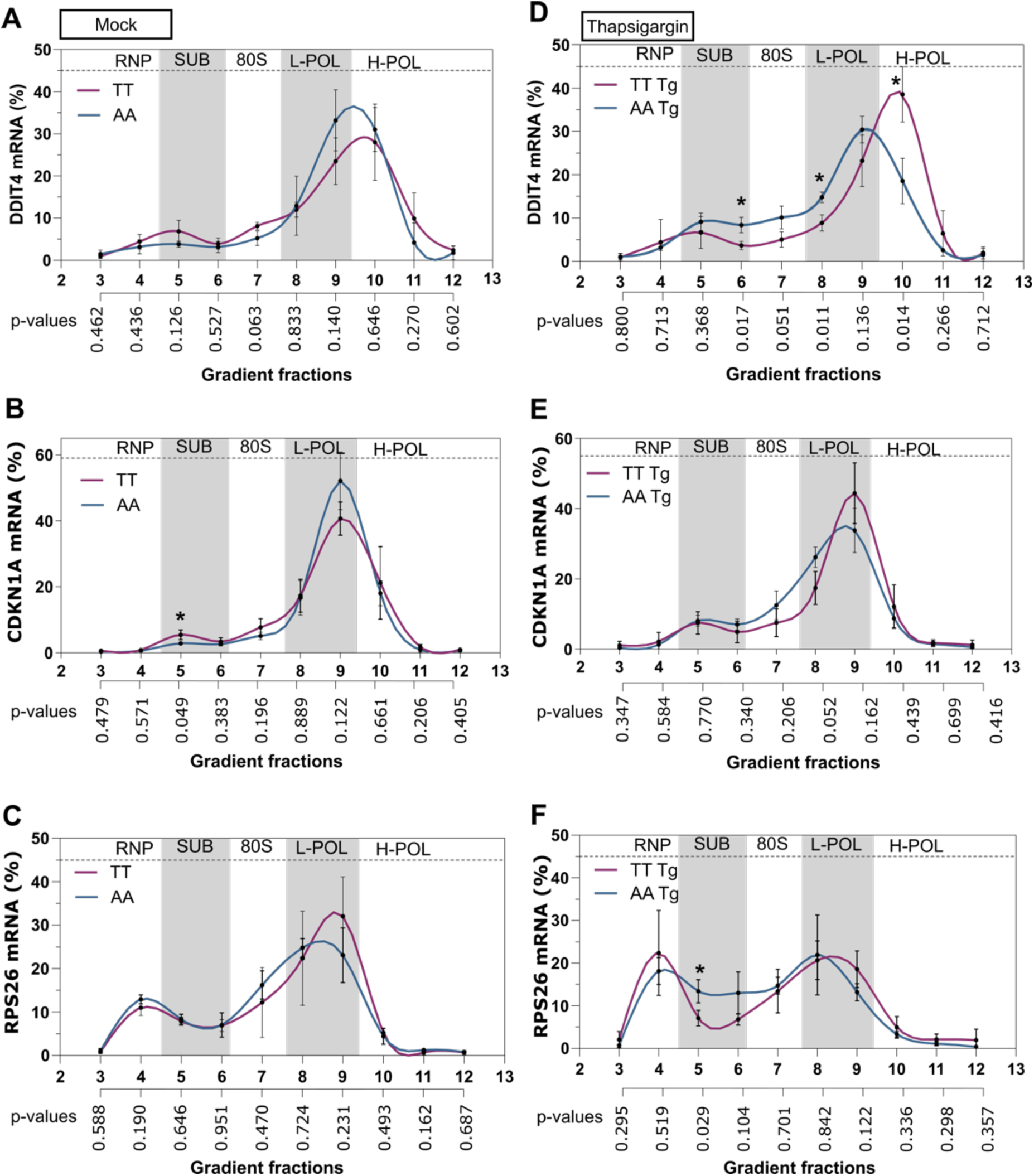
DDIT4 rs1053639 impacts its own mRNA distribution in polysomes. Cytoplasmic lysates from HCT116 rs1053639 homozygous (TT, AA) clones treated with thapsigargin 100nM or vehicle for four hours were subjected to fractionation through sucrose density gradients (15 to 50%). Polysome profiles are shown in Figure S4A. **A-D**) The distribution of DDIT4 mRNA (mock or thapsigargin conditions) across the ten gradient fractions was analyzed across the two genotypes by RT-qPCR. The relative mRNA percentage in each fraction was plotted. Three independent biological experiments were performed. Presented are the average proportion and the standard deviations of the three replicates. Statistically significant p values at the unpaired t-test are indicated by an asterisk. The p-values are shown below the images. The shading pattern in each image refers to the sucrose-gradient fractionations (Figure S4A). RNP = light fraction corresponding to free RNA and ribonucleoprotein complexes; SUB = ribosomal subunits; 80S = monosomes. Fractions corresponding to the light (L-POL) and heavy polysomes (H-POL) are also highlighted. CDKN1A (**B-E**) and RPS26 (**C-F**) mRNAs were used as control transcripts and were analyzed using the same methodology as described for the DDIT4 mRNA.

No differences were also apparent for the CDKN1A and RPS26 mRNAs, except for a slight increase in the proportion of CDKN1A in subpolysomal fraction 5 in TT cells (**Figure 3B, 3C**). When the same analysis was performed after thapsigargin treatment, a shift was visible for the DDIT4 mRNA in TT cells, with significant higher proportion in polysomal fraction ten and corresponding reduction in subpolysomal and light polysomal fractions, significant for fraction 5 and 8 (**Figure 3D**) Thapsigargin treatment led to a general reduction in the polysome association for the three mRNAs, which was more pronounced for the RPS26 mRNA (**Figure 3E, 3F**). A higher proportion of RPS6 mRNA in subpolysomal fraction 5 in AA cells was observed. Collectively, these results are consistent with the higher DDIT4 protein expression in TT cells, particularly in response to thapsigargin treatment.

### The rs1053639 tranSNP genotype impacts DDIT4 mRNA fate through the RBMX protein

We developed predictions for allele-specific binding starting from a list of 552 RNA binding proteins (RBPs) with consensus binding motifs (33), using the TESS software (DOI:10.1002/0471250953.bi0206s21) (34) (**Table S3**). The predictions and the allele-specific binding scores pointed to ACO1, YBX1, and particularly RBMX as RBPs that might differentially bind to the T/A alleles. Recent PAR-CLIP data in HEK293T cells (35) highlighted that RBMX binds in the 3’UTR of DDIT4 mRNA; therefore, we decided to focus on this target RBP. RBMX is mainly nuclear located, can interact with structural features of mRNAs, and was reported to modulate mRNA splicing, processing, and degradation (36–39). Hence, we performed RNA electromobility shift assays using the full-length recombinant RBMX protein and RNA probes mimicking the secondary structure of T and the A alleles of the endogenous DDIT4 mRNA. For the RNA probe design, we took advantage of three learning-based methods (EternaFold, CONTRAfold, and Ufold), two thermodynamic-based methods (ViennaRNA and RNAstructure), and one hybrid method (MXFold2) (40–43). Although the global structure of the 3’UTR was not significantly impacted, significant differences in the local secondary structure were observed after changing the base at the SNP site. RNA EMSA revealed that both RNA probes are bound by RBMX *in vitro*, with a possible preference for the T allele (**Figure 4A**). We further explored the publicly available web service AlphaFold 3 to generate accurate structure predictions for protein-RNA interactions. Interestingly, AlphaFold 3 provided several models confirming that the T allele RNA EMSA probe is located in a more favorable spatial conformation than the A allele probe for RBMX binding (**Figure S5A**). It also predicts that the C-terminal low complexity domain of RBMX binds to DDIT4 mRNA.

**Figure 4.**
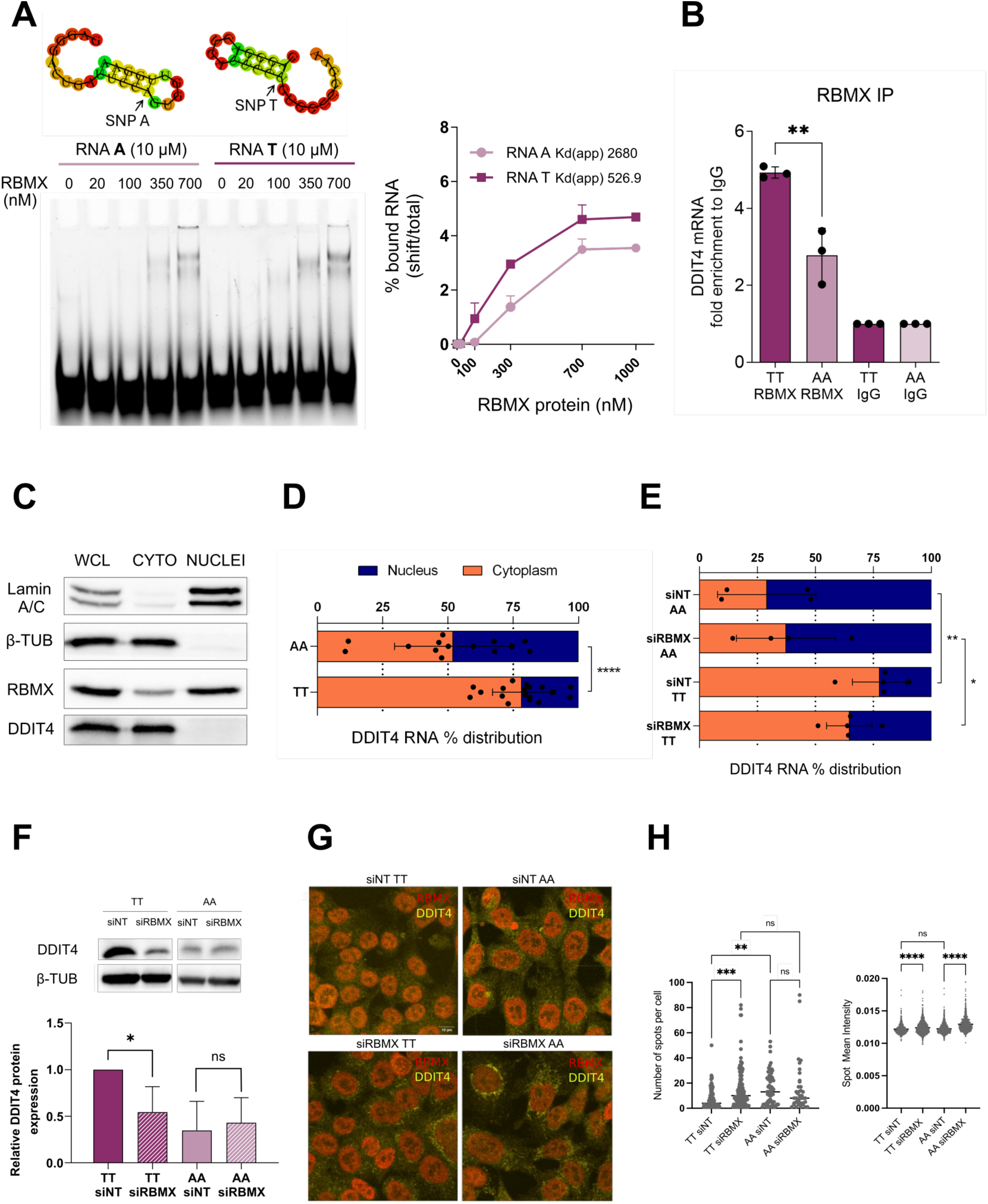
The rs1053639 tranSNP genotype impacts mRNA fate through the RBMX protein. **A**) **Left panel**, structural RNA probes were designed based on RNA folding prediction tools on the endogenous DDIT4 3’UTR sequence. RNA EMSA was performed using the full-length recombinant RBMX protein and the above-mentioned RNA probes. In the ‘0’ lane, the RNA probe was incubated with the binding buffer and the protein elution buffer. RNA-protein complexes were resolved on a native 6% gel. Right panel, quantification of the shifted bands in the REMSA experiments. Error bars plot the standard deviations among replicates. The apparent Kds obtained using non-linear regression fit are shown. **B**) The endogenous RBMX protein was immunoprecipitated from a whole cell lysate in TT and AA clones. Results are expressed as fold enrichment of the target DDIT4 mRNA relative to IgG. Three independent biological replicates were performed **p-value < 0.001, unpaired t-test. **C)** Cell fractionation was performed in HCT116 cells, and protein levels were evaluated. Lamin A/C (70-75 kDa) was used as a marker of the nuclear fraction, and β-Tubulin (55 kDa) was used as a marker of the cytoplasmic fraction. The DDIT4 protein was found exclusively in the cytoplasm. RBMX is mostly a nuclear protein, as previously reported. WCL: whole cell lysate. CYTO: cytoplasm. **D)** RNA levels were evaluated in TT and AA cells upon cell fractionation, followed by RT-qPCR. The distribution of DDIT4 mRNA is significantly different among TT and AA cells, with DDIT4 mRNA from TT cells being considerably more enriched in the cytoplasm. Data represent the mean ± SD of at least four independent experiments. The relative expression of the gene in the cytoplasm is calculated as the fold change of the Ct in the cytoplasm compared to the Ct in the nucleus, with nuclear and cytoplasmic RNAs eluted in the same volume of water. ****p-value < 0.0001. Unpaired t-test. **E)** DDIT4 mRNA levels in control (siNT) and in RBMX knocked-down (siRBMX) TT and AA cells were evaluated upon cell fractionation and RT-qPCR. The cytoplasmic localization of DDIT4 mRNAs was slightly reduced by RBMX depletion. Data represent the mean ± SD of three independent experiments. *p-value = 0.0114 **p-value = 0.0031, unpaired t-test. **F)** DDIT4 protein levels in control (siNT) and RBMX knocked-down (siRBMX) TT and AA cells were evaluated upon cell fractionation, followed by western blot. Representative blot of DDIT4 protein levels across TT and AA clones (Top). Densitometry analysis of the blots from several biological replicates shows that DDIT4 protein levels in TT clones decrease to the levels of the AA clones upon RBMX knockdown (Bottom). *p-value < 0.05, unpaired t-test. **G)** Confocal images of smiFISH-IF of a TT and an AA cell clone after control (siNT) or RBMX silencing for 72 hours. RBMX protein is shown in red, and DDIT4 mRNA is shown in yellow. Scale bar is 10 μm. **H)** At least three fields (15-20 cells per field) for each coverslip and each condition were imaged, and cytoplasmic spots were quantified using Cell-Profiler 4.0.7 (Broad Institute, Inc.) and segmented as objects with a typical diameter range of 10 to 30 pixels. (Left panel), number of DDIT4 mRNA spots per cell. (Right panel), spot mean intensity. ** p< 0.01; *** p<0.001; **** p<0.0001, ordinary one-way ANOVA. ns = not significant.

We then performed RIP assays to evaluate this interaction in a cellular context. The antibody used proved to be effective for the RIP protocol (**Figure S5B**). We first performed RIP in parental HCT116 cells, followed by Sanger sequencing of the input and IP samples. However, we could not appreciate quantitative differences in RBMX binding across the two alleles with this assay (**Figure S5C**). We next used both the rs1053639 TT and AA clones to perform RIP-qPCR. The qPCR analysis revealed that RBMX binds to DDIT4 mRNA slightly less than the positive control CSDE1, and we could confirm the preferential binding to the rs1053639 T allele (**Figure 4B**). Given the nuclear localization of RBMX and its binding to DDIT4 mRNA, we decided to examine the relative subcellular localization of DDIT4 mRNA alleles. Specifically, nuclear and cytoplasmic fractions were obtained and verified at both protein (**Figure 4C**) and RNA levels (**Figure S5F**). Results indicate that the DDIT4 mRNA shows higher cytoplasmic localization in the TT clones while being significantly more nuclear in the AA ones (**Figure 4D**). To shed light on the contribution of RBMX to this phenotype, we confirmed that our edited cells expressed it at comparable levels, and we used siRNAs to deplete its expression by ∼50% (**Figure S5D, S5E**). Nuclear and cytoplasmic fractions upon RBMX silencing were verified at the RNA level (**Figure S5G**). RBMX silencing led to a slight increase of nuclear DDIT4 mRNAs in TT clones (**Figure 4E**), and, importantly, to a significant decrease of DDIT4 protein levels (**Figure 4F**). No significant changes were observed in AA cells. Single-molecule FISH showed a punctuated distribution of the DDIT4 mRNA, particularly in the perinuclear region of the cytoplasm (**Figure 4G, S5H**). The number of granules per cell was higher in the AA compared with the TT clones (**Figure 4H**, left panel), while the spot mean intensity and area were not significantly different across clones (**Figure 4H**, right panel; **S5I**). Interestingly, the depletion of RBMX led to a significant increase in granularity in TT cells that reached the levels observed in the AA ones (**Figure 4G-H, S5I**). Hence, we suggest that RBMX contributes to the rs1053639 allele-specific regulation of DDIT4 protein levels through multiple mechanisms that include export, cytoplasmic localization, and translatability of the DDIT4 mRNA.

### The rs1053639 tranSNP genotype impacts on mTOR pathway markers

Since DDIT4 is an established inhibitor of mTORC1, (20,44,45) we next sought to investigate whether the effect of rs1053639 on DDIT4 protein levels would impact features of the mTOR pathway as well. In particular, we analyzed the levels of EIF4EBP1 and S6K phosphorylation both in basal culture conditions and in response to Thapsigargin treatment. We observed unexpectedly higher levels of relative phospho-eIF4EBP1 and phospho-S6K for TT clones in basal conditions, suggesting that our clones may have acquired compensatory mechanisms to balance their metabolism (**Figure 5A-D**) (46,47). Instead, in response to Thapsigargin treatment, TT clones were associated with a significant reduction in those mTORC1-dependent phosphorylations, which was not changed in the AA clones. While the AA clone #6 showed a distinct eIF4EBP1 phosphorylation pattern, analysis on additional AA clones revealed that this feature was not attributable to the AA genotype but only to this specific clone. Analysis of the protein levels of the ER stress response markers ATF4, a transcription factor regulating DDIT4 expression (48,49) (**Figure 5E**), and of the transcript levels of CHOP, spliced XBP1, and SESN2 (**Figure 5F**) did not reveal differences among HCT116 clones with the three rs1053639 genotypes in response to Thapsigargin (**Figure 5G**).

**Figure 5.**
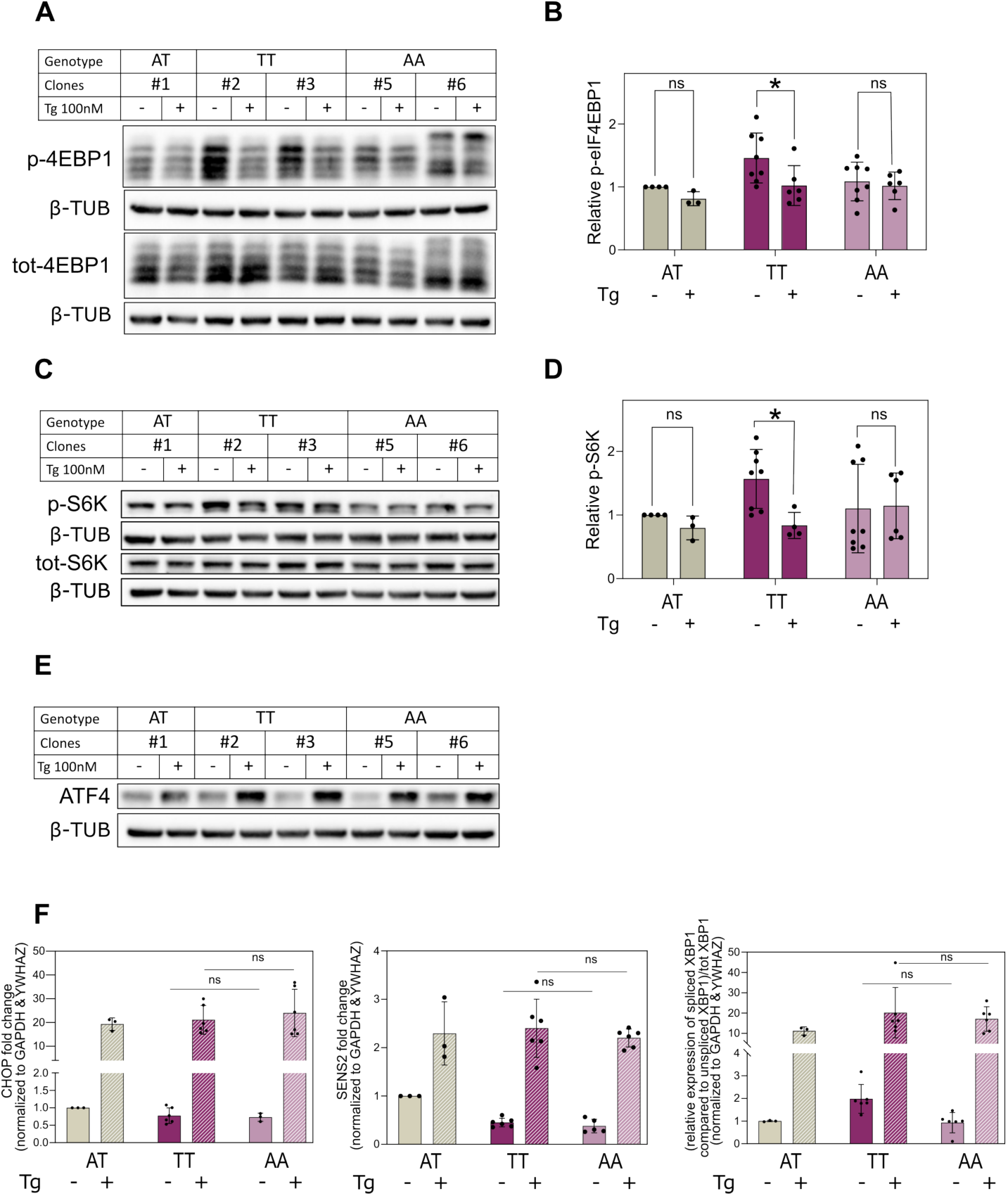
DDIT4 rs1053639 genotype-specific effect on the mTOR pathway. HCT116 parental cells and clones differing for the rs1053639 homozygous genotypes were treated with either thapsigargin 100nM (Tg) or vehicle control for 4 hours. **A, C**) Cell lysates were analyzed by immunoblotting for the phosphorylation state and levels of the mTORC1 target eIF4EBP1 and S6K proteins. **B, D**) Densitometry analysis of all blots reveals that significant inhibition of the phosphorylation of both eIF4EBP1 and S6K proteins is apparent only in the rs1053639 TT homozygous clones. *p-value <0.05. ns = not significant, unpaired t-test. **E, F**) ER stress response markers are activated in HCT116 cells in response to thapsigargin treatment independently of the rs1053639 genotype. **E)** Western blot analysis of ATF4 protein induction upon thapsigargin (Tg) treatment in HCT116 cells (A/T heterozygous) and DDIT4 rs1053639 homozygous clones (TT, AA). β-Tubulin was used as a reference. F) Relative CHOP, SENS2, and spliced XBP1/unspliced XBP1 mRNA expression as measured by RT-qPCR and normalized to GAPDH and YWHAZ ns = not significant, unpaired t-test.

We concluded that the rs1053639 SNP status does not broadly impact the activation of the ER-stress response upstream of DDIT4. Consistent with the effect of Thapsigargin treatment, a more pronounced reduction in phospho-eIF4EBP1 was observed for TT homozygous cells also in response to Nutlin treatment (**Figure S3A, D**).

Next, we investigated whether the differences in the mTOR pathway led to changes in global translation or proliferation of the HCT116 edited clones. The global translation was monitored either in mock culture conditions or in response to Thapsigargin treatment using a quantitative analysis of the Translation Efficiency (TE) measured as the relative abundance of polysomes compared to the 80S from polysomal profiling or the measurement of newly synthesized peptides using either a commercially available global protein synthesis kit or a puromycilation assay (**Figure 6A-C**, **Figure S4A**).

**Figure 6.**
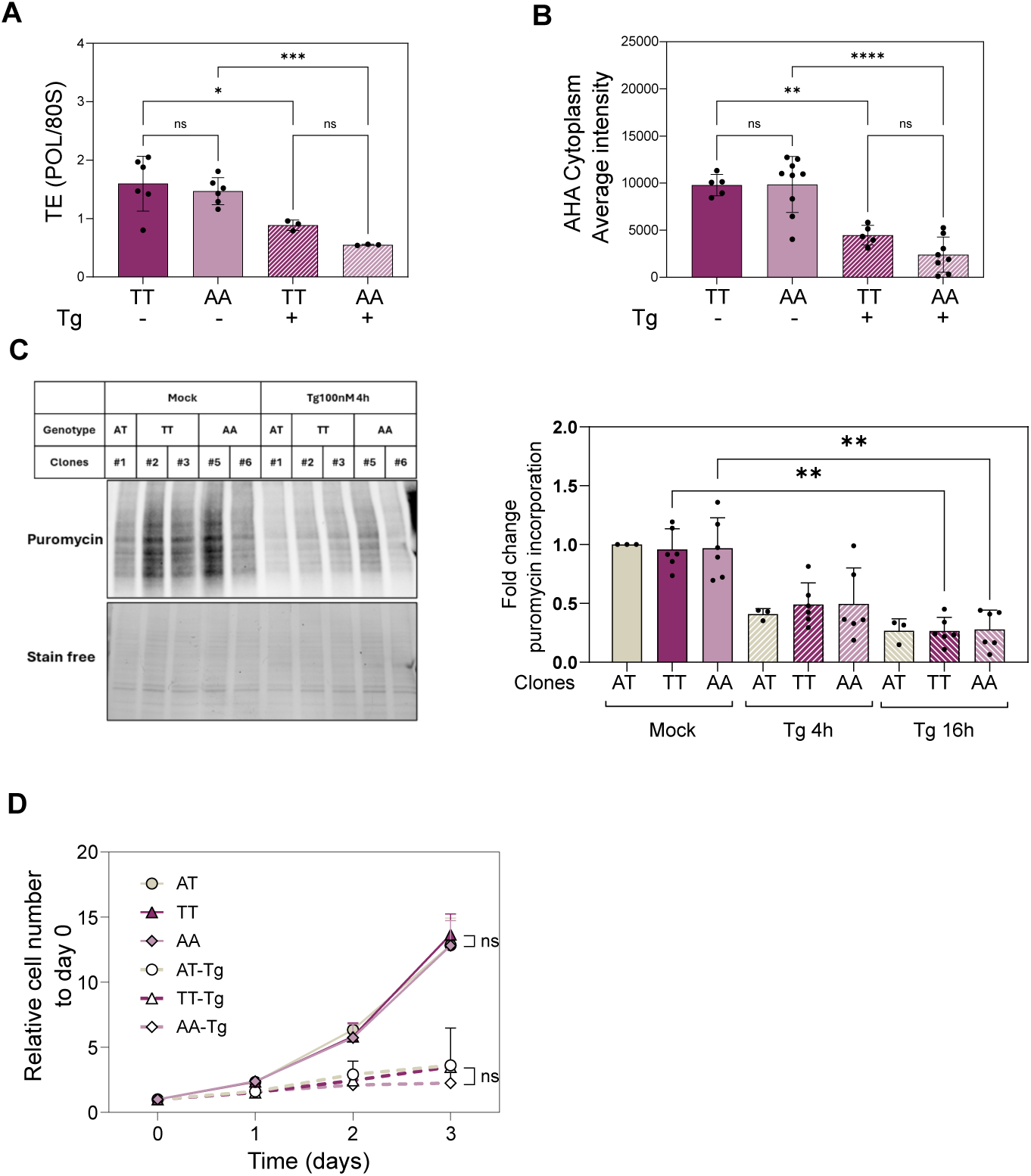
The DDIT4 rs1053639 genotype does not significantly impact global translation and cell proliferation in response to Thapsigargin. **A**) Global Translation Efficiency (TE) measured from polysome profiles (see Figure S4) prepared using sucrose gradient fractionation of homozygous TT and AA HCT116 cells treated as in Figure 1A. **B**) Global translation was measured using a Click-it reaction assay after incubation of the cells with modified methionine for 30 minutes. Homozygous TT and AA HCT116 cells grown in mock conditions or treated with 100 nM Thapsigargin for four hours were tested. In both assays, Thapsigargin led to a reduction in global translation. **C**) Global translation measured by a puromycilation assay. A representative Western blot is presented on the left: immunodetection with an antibody targeting puromycin, and the loading control is shown. The bar graph on the right plots the results of the densitometric analysis of the blots plotted as fold change of puromycin incorporation for the indicated HCT116 clones and treatment conditions. Average, standard deviations, and individual replicates are shown. *P-value < 0.05; **P-value < 0.001; ***P-value < 0.0005. **** P-value < 0.0001, two-tailed, unpaired t-test. **D**) HCT116 cells parental and clones differing for the rs1053639 homozygous genotypes were treated with thapsigargin 10nM of vehicle control for 72 hours. The proliferation of the clones was followed every 24 hours by the Operetta HCS system. Relative cell number curves were plotted as the mean of several replicates ± standard deviation (SD). ns, not significant. Ordinary two-way ANOVA.

In all cases, there were no significant differences in global translation comparing TT and AA rs1053639 clones, although the latter showed a trend for a stronger reduction in the perturbed conditions. The relative proliferation of the clones was similar in the mock condition and in response to thapsigargin (**Figure 6D**), while TT clones showed slightly higher resistance to Nutlin treatment (**Figure S4B**). We concluded that the rs1053639 tranSNP genotype did not show an overt difference in global translation or cell proliferation.

### The rs1053639 tranSNP AA genotype shows a competitive growth advantage in prolonged culture conditions

Given the contribution of DDIT4 in the negative modulation of a critical and intricate metabolic hub such as mTOR (50), we tried to tease out possible differences in cellular phenotypes due to the rs1053639 genotypes through co-culture experiments where cells could experience fluctuations in nutrient availability in the media. To visualize and count the contribution of each genotype to the total co-culture population, derivative GFP or RFP positive clones were obtained with high clonal purity. Cells were seeded in 6-well plates, and relative cell number was measured either at Operetta High Content Screening (HCS) or by FACS, examining the population every four days, up to fifteen days. Results were expressed as relative fitness, as recently described (51) (**Figure 7A**). Interestingly, while the rs1053639 TT cells showed higher fitness at early times, AA cells progressively gained a proliferative advantage. This result was not influenced by the presence or the type of fluorescent marker (**Figure S6A**). Notably, the depletion of RBMX both in TT and AA cells three days prior to the co-culturing reversed the proliferative advantage phenotype of the AA cells (**Figure 7B**). To further explore the significant difference related to rs1053639, cells were sorted seven days after co-culture, and the two rs1053639 genotypes were examined separately. No significant difference in cell cycle distribution was observed (**Figure 7C**). Finally, the co-culture experiment was repeated in the presence of the Q-VD-OPh pan-caspase inhibitor (52). The progressive advantage of the AA genotype was confirmed, suggesting that apoptosis was not involved in this phenotype (**Figure 7D**). Doxorubicin 3µM was included as a control to induce apoptosis (53,54) (**Figure S6C, S6D**).

**Figure 7.**
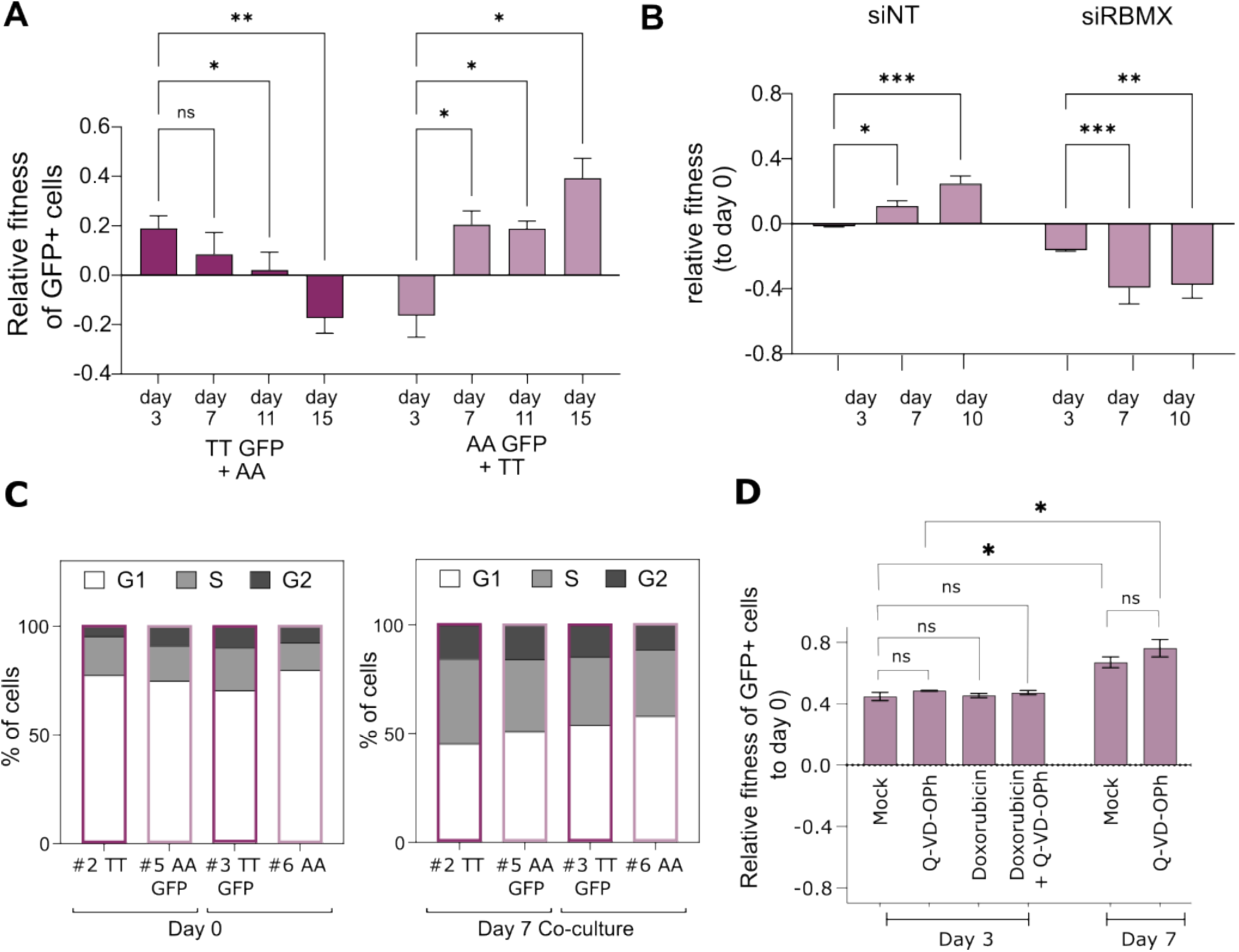
DDIT4 rs1053639 impacts cellular dynamics and survival under stress and competitive conditions. **A**) DDIT4 clones harboring the TT genotype were transduced with Lentiviral particles to express EGFP constitutively. After transduction, cells were treated with 5 µg/ml blasticidin for at least 15 days, to reach at least 95% of GFP+ cell populations. DDIT4 clones harboring the AA genotype (not fluorescent) were mixed ∼1:1 with EGFP-expressing TT genotype cells and co-cultured for 15 days. The relative growth was assessed by flow cytometry, analyzing 10K cells. Cellular fitness of the GFP+ clones relative to GFP-competitor clones was calculated as follows: pGFP+(t) = NGFP+(t) / (NGFP+(t) + N GFP-(t)), W = ln ((pGFP+(ti) /pGFP-(ti) ) / (pGFP+(t0) /pGFP-(t0)), adapting a described method (51). The error bars represent the mean ± SD of at least eight independent experiments (Four TT GFP+, four AA GFP+). *P-value < 0.05; **P-value <0.01, ordinary two-way ANOVA. **B)** Relative fitness of rs1053639 AA homozygous cells (GFP labeled) in co-culture experiments with the TT homozygous counterpart. Both TT and AA-GFP cells were transfected to deplete RBMX (siRBMX) (or control, siNT) 3 days before starting the co-culture and silenced again at day 3 and day 7 during the co-culture experiment. The relative fitness of GFP-positive cells is shown. *P-value < 0.05; **P-value <0.01, ***P-value <0.001, ordinary two-way ANOVA. **C**) Cell cycle analysis by flow cytometry (FACS) for single clones before co-culture (Left) and sorted clones seven days after co-culture (Right). **D**) TT and AA clones were co-cultured as described in A) up to seven days, in the presence of the pan-caspase inhibitor Q-VD-OPh (10 µM) or vehicle control, and analyzed by FACS at days 0, 3, and 7. As a control, co-cultured cells were treated with Doxorubicin (3µM) +/- Q-VD-OPh (10 µM) for 3 days. The relative fitness of GFP-positive cells across time points and conditions is shown. *P-value < 0.05, two-tailed unpaired t-test.

### The rs1053639 tranSNP AA genotype showed faster growth and a competitive advantage when xenografted in Zebrafish embryos

The proliferation experiments in co-culture suggested to us that the rs1053639 tranSNP genotype could impact cell growth in challenging environments where nutrient concentrations vary, and different cell types can compete. To explore this hypothesis and also gain some information on the potential aggressiveness of our edited cancer cell lines, we developed xenograft experiments in Casper Zebrafish embryos (55), following a published protocol adjusted for our experiments (56). An equal number of cells of the two different genotypes labeled either with GFP or RFP, as described earlier, were injected into the swim bladder of three-day post-fertilization embryos. Cells were then imaged the following day and again 48 hours later. Results, plotted as a percentage of tumor cells’ growth comparing 24 to 72 hours post-injection, showed that, when cells of the two genotypes were pre-co-cultured *in vitro* for three days prior to injection in the embryos, the growth advantage of the AA genotype was again apparent (**Figure 8A, B**, **S7A**).

**Figure 8.**
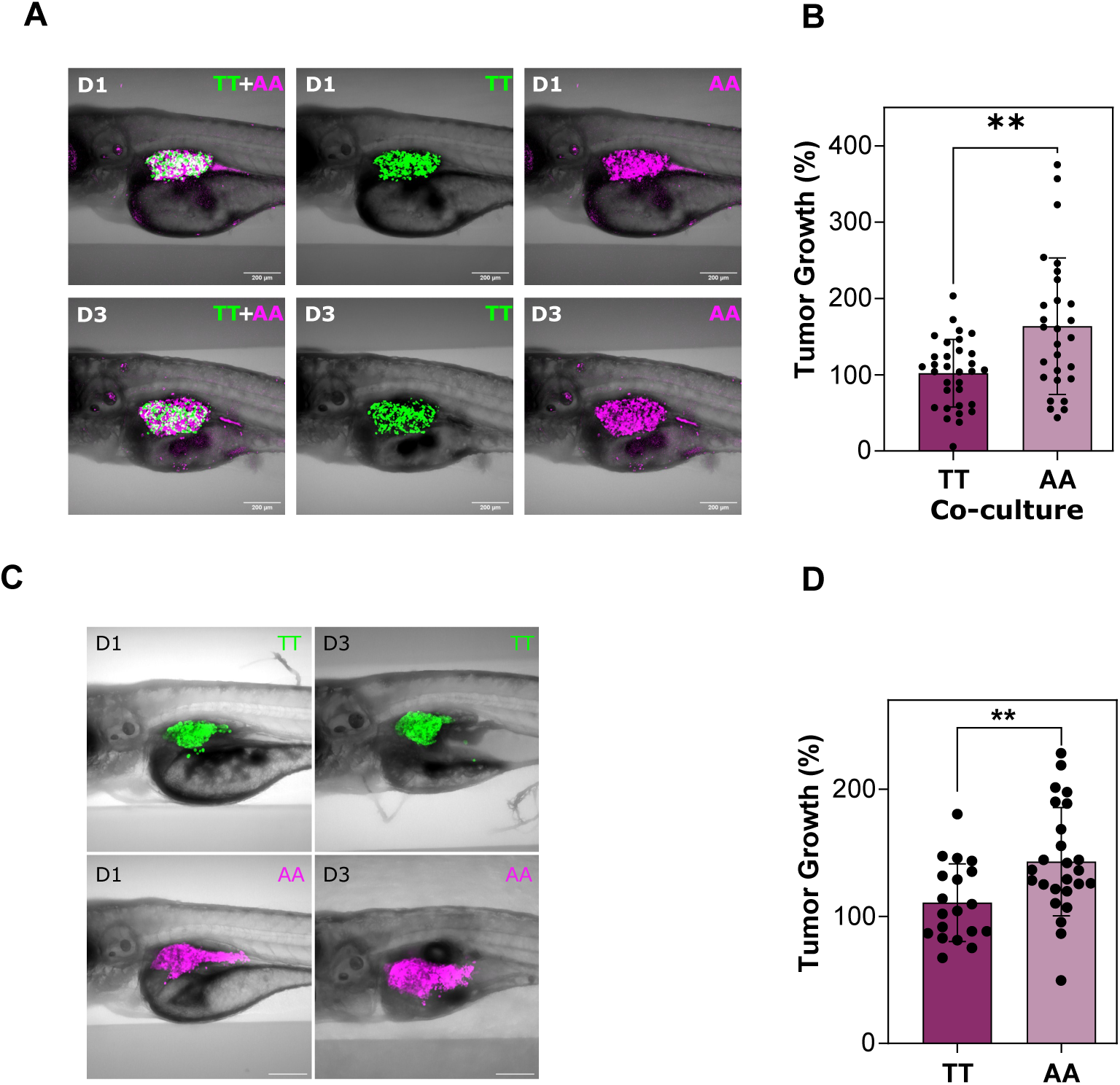
DDIT4 rs1053639 impacts tumor growth and genotypic competition in zebrafish larvae xenotransplants. DDIT4 rs1053639 homozygous clones, transduced with lentiviral particles to constitutively express either EGFP or MiRFP as described in Figure 5A, were xenotransplanted into zebrafish larvae two days post-fertilization. Tumor growth was monitored over a three-day period, with a minimum of 30 larvae per genotype condition. Representative confocal microscopy images were captured on day 1 and day 3 post-transplantation, with TT clones in green and AA in purple. **A**) DDIT4 clones expressing GFP (TT clones) were mixed∼ 1:1 with RFP-expressing clones (AA genotype). They were initially co-cultured *in vitro* for four days, followed by xenotransplantation into ∼30 zebrafish larvae for an additional three days of *in vivo* co-growth before confocal imaging. **B**) Bar graph illustrating the percentage of tumor growth relative to day 0 of transplantation, with each data point representing an individual larva. AA clones harbor significantly higher fitness. **P-value < 0.01. paired t-test. **C**) TT and AA clones were separately xenotransplanted into zebrafish larvae, which were followed for three days and imaged by confocal microscopy. **D)** Quantitative analysis of tumor growth indicates a significantly faster growth rate for AA than TT cells *in vivo*. Each data dot represents one larva. **P-value < 0.01. paired t-test.

Furthermore, we evaluated the proliferative potential of the TT and AA cells separately in the *in vivo* setting. Notably, the AA cells grew more compared to the TT ones as well (**Figure 8C, D, S7B**). Collectively, co-culture experiments indicate that rs1053639 AA cells display a competitive advantage over TT cells both *in vivo* and *in vitro*, although the underlying mechanisms remain to be clarified, and it does not seem to fit under a classical cell competition response (57).

### The rs1053639 tranSNP status can have prognostic significance in cancer patients

The phenotypic differences related to the rs1053639 genotype, a relatively common SNP in the human population, led us to explore whether this tranSNP status can impact clinically relevant variables in cancer patients. Based on the results obtained with the three genotypes, particularly the reduced control of the mTOR pathway of the AA homozygous cells, we decided to focus on a recessive model, comparing AA homozygous patients with AT plus TT. Using TCGA data we observed worse prognosis for AA patients in both colorectal and esophageal cancers (**Figure S8A, S8B**). DDIT4 mRNA expression showed a prognostic value with high expression, which is associated with reduced survival probability in lung adenocarcinoma but higher survival probability in esophageal cancers, according to recent reports (58–60). In colon adenocarcinomas, DDIT4 mRNA expression is significantly higher than healthy tissues (p-value 8.9e^-21^) (**Table S8**). Further, data in DepMap indicate a reasonable level of correlation between DDIT4 protein and mRNA levels (Pearson 0.517, linregress p-value 8.82e^-5^) (**Figure S8C**). We began exploring the correlation between rs1053639 genotype and DDIT4 protein levels by IHC in seven biopsies of primary colorectal adenocarcinoma and adjacent resected normal tissues. To this aim, DNA from peripheral blood was obtained from the same group of samples, and the DDIT4 SNP status was genotyped (**Figure S8D**). The samples were classified by certified pathologists in two categories based on IHC intensity. The average signal intensity on the different samples, first classified as Low, Intermediate, and High, was re-evaluated using Fiji, and it was found that the average intensity of the Intermediate samples was comparable to the average intensity of the Low samples. Therefore, it was decided to reduce the classes from three to two (Low and High). Overall, cytoplasmic expression of DDIT4 protein was higher in the cancer samples than in healthy tissue from the same patient. Only one sample (TT) received a high score both for normal and tumor sections (**Figure S8D**, **E1**, and **E2**). Further studies are needed to establish the value of rs1053639 SNP status as a prognostic marker.

## Discussion

### Variation in allelic imbalance in polysomal RNA as a tool to identify functionally relevant SNPs

Our approach to identifying tranSNPs is based on computing and comparing relative changes in allele counts across mRNA fractions, building upon the methodology previously introduced (9), and the underlying assumption in our approach is that the interaction of an mRNA with polysomes is directly proportional to the amount of protein being produced (61). By fully leveraging biological replicates, our enhanced method eliminates the need for a predefined threshold to detect allele-specific expression. Instead, it evaluates differential allelic fractions between RNA-seq data from the two distinct cellular compartments using pairwise statistics, thereby improving the robustness of the analysis. Despite the limitations imposed by low coverage, which introduced noise in allelic fraction estimates and constrained our ability to detect and characterize SNPs with significant allelic imbalances, our findings support the presence of signal in over 11% of analyzable genes. Each of these genes harbors one or multiple imbalanced SNPs, collectively accounting for approximately 7% of all analyzable SNPs, aligning with previous reports (8,9).

### Mechanism of rs1053639 impact on DDIT4 protein expression

For rs1053639 A/T we observed a significant reduction in the alternative allele (A) in polysome-associated mRNAs in the mock condition. Using edited HCT116 clones homozygous for the alternative and reference alleles, respectively, we observed a marked difference in DDIT4 protein expression associated with the different SNP status. In our analysis pipeline, we also used predictions of trans factors binding to mRNAs. In this respect, rs1053639 is predicted to affect an RBMX binding motif. Given that RBMX is reported as prevalently nuclear (36), an observation confirmed also for HCT116 cells (**Figure 4C**), we measured DDIT4 mRNA localization comparing the TT and AA homozygous clones. In general, RNA binding proteins binding 3’-UTR sequences can contribute to subcellular mRNA localization and mRNA fate (68). Results indicated higher cytoplasmic localization for the T allele. The transient silencing of RBMX slightly increased the nuclear localization of the T allele and significantly increased the number of perinuclear granules of DDIT4 mRNA. This effect was associated with a significant reduction of DDIT4 protein expression with the TT genotype, making it comparable to the levels of the AA clones. In cells of the AA genotype, the number of DDIT4 mRNA perinuclear granules was lower than in TT cells in the mock condition. Interestingly, RBMX depletion did not significantly change the nuclear localization of the DDIT4 mRNA A allele, the number of perinuclear granules per cell, or DDIT4 protein levels. The observed increase of perinuclear DDIT4 mRNA granules in the AA clones, irrespective of RBMX expression, and in the TT clones only upon RBMX depletion, suggests an impact of the tranSNP, together with this trans factor, on mRNA export and translation. Indeed, mRNA granules near the nuclear membrane have been considered intermediate sites during mRNA export (69–71). Moreover, recent reports highlighted the relevance of mRNA localization in cytoplasmic compartments for controlling its protein production (68).

Collectively, our validation experiments confirmed the observation of allelic imbalance in polysomes and provided evidence that rs1053639 can impact DDIT4 protein expression through a mechanism that comprises mRNA localization, export, modification, and translation efficiency.

### Implications of rs1053639 allelic status on DDIT4-mediated functions

DDIT4 is an established important negative modulator of the mTOR pathway, acting at the level of mTORC1. Consistently, its gene expression is modulated in response to ER stress, with ATF4 as an important transcription factor. It is also inducible by hypoxia and is considered a p53 target gene, although recent data proposed that the contribution of p53 is indirect, via the transcription factor RFX7 (20). In general, mTOR inhibition is growing in relevance in p53-dependent tumor suppression (72). The negative modulation of mTORC1 by DDIT4 is rather established, although reports differ on the exact mechanism (45,50) and its relative relevance in different cell types (73). DDIT4, as the extended name testifies, is considered a player in the DNA damage response, although also, in this regard, mechanistic details are missing. Overall, DDIT4 function in cancer is considered to be tumor suppressive. For example, a relatively recent screen indicated DDIT4 in a short list of p53 pathway targets with strong tumor suppressive potential. In agreement with the centrality of the mTOR pathway for general cell metabolism, DDIT4 has emerged as an important gene in contexts other than cancer, such as liver disease, neurodegeneration, pulmonary injury, and retinopathy (64,74–76). In fact, post-transcriptional regulation of DDIT4 expression has been the subject of different studies that can bear relevance for understanding the impact of rs1053639. For example, two studies pointed to the important role of an antisense transcript (DDIT4-AS1) corresponding to a large portion of DDIT4 3’UTR and of the 3’ coding sequence, although with different conclusions on the role of the antisense transcript on the sense one (27,28). rs1053639 is included in the antisense. We checked if the allelic status could somehow impact the amount of spliced DDIT4-AS1. In HCT116, the antisense was expressed at a very low level, and there were no appreciable differences among the edited clones (**Figure 2**), indicating that it does not contribute to the difference in DDIT4 protein expression in our context.

In HCT116 cells, our experiments demonstrate that rs1053639 SNP status significantly impacts DDIT4 protein levels. The lower expression of DDIT4 in AA homozygous clones is even more pronounced when the gene transcription is induced by ER stress or by Nutlin. Hence, the tranSNP we identified is predicted to impact p53-dependent and, more in general, stress-dependent modulation of DDIT4 with potentially appreciable consequences on the negative modulation of mTOR, impacting translation control, proliferation, autophagy, and cell survival. Given the complex regulation of the mTOR pathway (77), the impact of DDIT4 is expected to be at least in part influenced by tissue type. In HCT116 cells, higher DDIT4 protein levels in the TT homozygous clones are associated with stronger repression of mTORC1 markers. Despite this, TT and AA clones showed similar proliferation in culture plates. However, in vitro co-culture experiments showed a higher proliferation of AA cells. The two separated populations did not show evidence of cell cycle checkpoint activation or apoptosis induction. Silencing of RBMX, which negatively impacts DDIT4 protein levels in TT but not AA homozygous cells, led to a reversal of that proliferation advantage. The results of the xenograft transplantation experiments in Zebrafish embryos were also consistent with higher proliferation of HCT116 cells homozygous for the rs1053639 A allele.

In general terms, DDIT4 is considered a tumor suppressor gene. However, gene expression data from TCGA indicate an almost equal number of cancer types where DDIT4 expression is higher in normal tissues than in cancer compared to the opposite trend. Also, in terms of survival analysis, high DDIT4 expression has been associated either with a better or worse prognosis. Although the data is exploratory in nature, we found consistent prognostic predictions when TCGA patients’ cohorts were stratified based on DDIT4 mRNA levels or for the rs1053639 genotype for LUAD, ESCA, and COAD.

### Study Limitations

The limited coverage of our RNA-seq dataset resulted in a relatively small number of heterozygous SNPs available for analysis. This was further constrained by our conservative approach, which requires the heterozygous state of an mRNA variant, a minimum of twenty reads, and consistency among four biological replicates of total and polysomal RNA sequencing that are then compared to nominate a tranSNP candidate. This strategy reduces the number of genetic variants that can be analyzed with a chosen cell line but enhances the likelihood of detecting true signals. Validation experiments using orthogonal approaches and reporter assays were limited to four selected candidates of which three showed consistent results across methods. While the heterozygous state is a requirement to start our analysis, the phenotypic impact of a variant is expected to be most pronounced in homozygous states. Therefore, confirming the causal role of a tranSNP on cell phenotypes requires the development of genomically edited, isogenic cell clones differing for SNP status, a time-intensive process that, in our study, was performed only in HCT116 cells.

## Conclusions

We have developed a robust pipeline to catalog genetic variants associated with allele-specific changes in mRNA translation, leveraging biallelic mRNAs and polysomal profiling as a proxy for translation efficiency (**Figure 9**).

**Figure 9.**
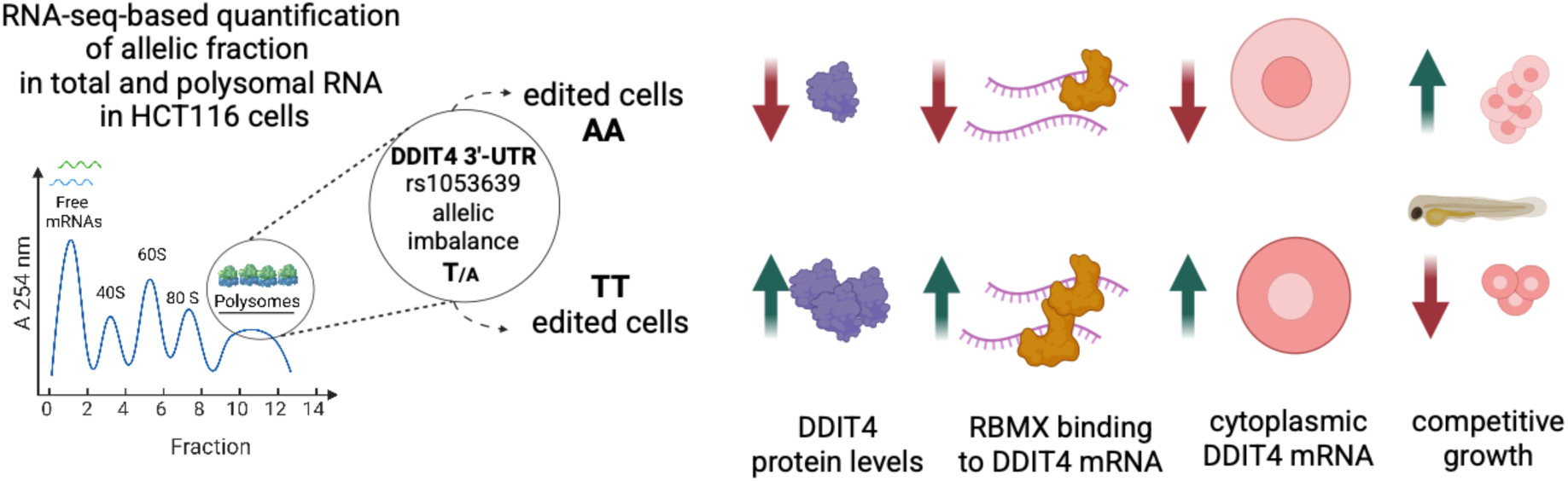
Graphical Summary. Scheme of the approach taken to identify tranSNPs and of the results obtained with rs1053639 in DDIT4 with the HCT116 cell model.

Using this approach, we identified 40 SNPs that exhibit allelic imbalance when comparing total and polysomal RNA. Focusing on rs1053639 in the DDIT4 3’UTR, we found that the alternative (A) allele is associated with reduced DDIT4 protein levels in the human colon adenocarcinoma-derived HCT116 cells. This reduction correlates with a higher abundance of the A allele in the nucleus and an increased number of DDIT4 perinuclear cytoplasmic mRNA granules. These allele-specific differences are mediated by the RNA-binding protein RBMX. Cells homozygous for the alternative A allele exhibit lower DDIT4 protein levels, leading to diminished negative regulation of the mTORC1 pathway, enhanced growth potential in zebrafish xenografts, and poorer prognosis in selected TCGA cancer cohorts. More broadly, our approach provides a powerful framework for identifying genetic drivers of inter-individual variability in mRNA translation, shedding light on mechanisms that can influence cancer progression.

## Materials and methods

### Cell lines and cell culturing

The human cell line HCT116 was purchased from the Biological Resource Center ICLC Cell bank. Cells were cultured in standard Roswell Park Memorial Institute (RPMI) 1640 medium (GIBCO) supplemented with 10% fetal bovine serum (FBS), 100 units/ml penicillin, 100 mg/ml streptomycin antibiotic mix (GIBCO), and 2 mM Glutamine (GIBCO). The cells were maintained in a humidified atmosphere at 37°C and 5% CO_2_.

### Dataset and data preprocessing

The RNA-sequencing data of RNA extracted from polled polysomal and total cytoplasmic fractions of HCT116 cells cultured either in mock condition or treated with the MDM2 inhibitor Nutlin was produced in our earlier study (13). The raw data is deposited in GEO, under the GSE95024 accession number. Single-end RNA-seq data was processed to obtain from row FASTQ files properly processed BAM files. The trimming step, essential to remove sequence adapters, was performed with Trimmomatic (78). The data were aligned with STAR (79) to the reference genome GRCh38 to obtain a BAM file for each sample. To improve the characterization of SNP positions, a step of deduplication to remove PCR artifacts was performed by employing PICARD module MarkDuplicates (broadinstitute.github.io/picard/). Then, GATK (80) was employed to split, realign, and recalibrate the data in order to improve the quality of the alignment. The pileup of SNPs positions was performed with ASEQ (81) using parameters mode=0, mbq=20, and mrq=20 in order to obtain allelic fraction (AF) calculations. Only SNPs in exonic and UTR regions extracted from dbSNP were considered. We emphasize that, as expected, the use of single-end RNA-seq data led to high duplication rates (on average 70%), limiting the number of SNPs we could characterize and analyze. Nevertheless, this step, along with subsequent quality control processes, was essential to ensuring high-quality AF estimations.

### Differential ASE analysis

The allelic fraction at SNPs’ positions was used to compute differential allele-specific expression (ASE) events between total and polysomal RNA fractions. In order to capture those events, we started by identifying all SNPs that were heterozygous in at least one biological replicate of the two fractions. More specifically, to define the analyzable SNPs, for the data obtained both in mock and Nutlin-treatment conditions, we searched for SNPs having a value of AF between 0.2 and 0.8 (i.e., heterozygous genotype) in at least one replicate of one of the two fractions (polysomal or total) and that were confidently observable, i.e., with a depth of coverage of at least 20x of the SNP site in the each fraction, condition, and biological replicate. A total of 1060 SNP/condition pairs were identified. Of these, only those with AF measures available across all four biological replicates and both fractions were then considered for differential ASE analysis (N=582, Table S1).

To assess the presence of a differential ASE event at a SNP position we extended the methodology we developed (9). In particular, for each SNP we exploited the concordance of differential AF measures across pairs of polysomal and total RNA biological replicates by performing paired t-test statistics. Although none of the 41 SNP/condition pairs with nominal p-value ≤ 0.05 kept significance after FDR correction, likely due to the low coverage that increased the noise in AF estimations, overall, 11.8% of the analyzed genes had one or multiple SNPs (in LD) with a p-value ≤ 0.05, strongly supporting the presence of a true signal. A list of SNPs with nominal p-values ≤ 0.05, denoted as putative tranSNPs, were hence retained for downstream analysis and experimental validation.

### Expression analysis of DDIT4 in cancer and normal tissue samples

To test whether the gene containing the rs1053639 tranSNP exhibits differential expression in tumor versus normal tissue samples, we compared transcript expression values (normalized RPKM values) of the DDIT4 gene exploiting primary tumor and normal tissue samples from the TCGA COAD dataset. In detail, we downloaded the gene counts for all genes from the recount 3 database (82), and RPKM values were computed using the function *getRPKM* from the recount 2 R package. Then, protein-coding transcripts were selected, and their RPKM values were normalized across samples using the R limma quantile normalization function. Then, focusing on the DDIT4 gene, the distributions of log2(RPKM+1) values across tumor and normal samples were compared using Wilcoxon’s test statistics, and p-values were finally corrected for multiple hypotheses using the FDR method.

### Survival assay using TCGA data

Genotype and survival data for TCGA patients were collected. Specifically, genotype information for rs1053639 tranSNPs was retrieved from (83), while survival data across different endpoints (Overall Survival, Disease-Specific Survival, Disease-Free Interval, and Progression-Free Interval) was retrieved from (84). A recessive model (AA+AB vs. BB) was built, and for each endpoint, a Kaplan-Meier survival curve was calculated along with summary statistics obtained from a multivariate Cox proportional hazards regression model, where age at diagnosis was included as a covariate. For each endpoint, the analysis was performed across 23 different cancer types, and p-values were corrected for multiple hypothesis testing, retaining only associations with a p-value < 0.05 and an FDR < 0.25. Survival analyses were performed using the survival R package.

### Identification of SNPs disrupting putative RBP consensus motifs

To investigate the RNA protein binding landscape at the rs1053639 tranSNPs locus, we retrieved 552 human RNA binding consensus motifs from the RBPDB database (33) and created a FASTA file containing a list of 60bp DNA sequences containing the reference and the alternative bases. Then, TESS software (34) was run to compute RBP consensus motifs scores. A set of log-likelihood-ratio-based scores were computed: the log-odds ratio of the match for the alternative and reference bases (*altScore* and *refScore*), and the maximum possible log-odds ratio for a match for a given RBP consensus motif (*maxScore*). The scores were filtered by keeping only the forward strand analysis. Results were retained only if the score for the reference or the alternative allele was at least 3 and greater than 0.5 when normalized by the *maxScore*.

### Polysome profiling, RNA extraction, and Sanger sequencing

HCT116 cells were grown in 150 mm dishes to 80% confluence followed by 100 nM Thapsigargin or vehicle treatment for 4 hours. Cells were then incubated for 10 min with 50 µg /ml cycloheximide at 37°C to disrupt the protein synthesis process and halt translation elongation, thereby immobilizing the ribosomes on the mRNAs and preventing ribosome disassembly. Cells were rinsed with ice-cold PBS containing 50 µg /ml cycloheximide and then gently scraped using 600 µl of ice-cold lysis buffer (20 mM Tris-HCl pH 7.5, 100 mM KCl, 5 mM MgCl_2_, 0.5 % Nonidet P-40, 0.2 U/µl RNasin® Ribonuclease Inhibitor [Promega], cOmplete™, Mini Protease Inhibitor Cocktail 1X [Roche], 100 µg /ml cycloheximide) and collected in an Eppendorf tube. The cells were then incubated on ice for 10 min and centrifuged for 10 min at 12,000 *x g* at 4 °C to remove the nuclei. The cytoplasmic lysate was then separated by ultracentrifugation in a Beckman SW41 rotor at 40.000 rpm for 1 hour and 40 minutes at 4°C after being loaded onto a 15-50% linear sucrose gradient dissolved in salt buffer (100 mM NaCl, 20 mM Tris-HCl pH7.5, and 5 mM MgCl2). The sucrose gradients were fractionated, and the absorbance was measured at 254 nm by the Teledyne Isco model 160 gradient analyzer equipped with a UA-6 UV/VIS detector. Based on density separation, polysomes were isolated from the fractions, and RNA was harvested from these polysomes with the TRIzol® reagent (Invitrogen ^TM^) following the manufacturer’s protocol. The RNA was then retro-transcribed, and the sequences containing the candidate TranSNPs were amplified by PCR using AmpliTaq Gold 360 Master Mix (ThermoFisher Scientific) and specific primers (**Table S4**). The PCR products were purified by the spinNAker GEL&PCR DNA purification kit (Euroclone) and subjected to Sanger sequencing by Mix2Seq Kit (Eurofins Genomics) following the manufacturer’s protocol. Total RNA was also harvested in parallel for Sanger sequencing. Analysis of the electropherograms was performed by Fiji software. Quantification was performed by assessing the area under the peaks of the tranSNP nucleotides as a metric to calculate allelic fraction. The normalization process considered the relative efficiency of the four nucleotides’ incorporation during sequencing, using the information of local surroundings. The allelic fraction for targeted genes was determined for at least three independent experiments. Student’s t-test was used to calculate the p-value.

### Targeted RNA sequencing

Targeted RNA-seq libraries were prepared using custom multiplex primers, each containing an ∼25-nucleotide (25-nt) gene-specific sequence and either a 33-nt Illumina adaptor (5′-TCGTCGGCAGCGTCAGATGTGTATAAGAGACAG-3′) at the 5′ end of the forward primer or a 34-nt Illumina adaptor (5′-GTCTCGTGGGCTCGGAGATGTGTATAAGAGACAG-3′) at the 5′ end of the reverse primer. Targeted RNA-seq libraries were prepared from DNase-treated RNA. cDNA was synthesized using a dTVN primer and purified with SPRIselect beads to remove excess primers. Fifty nanograms (50 ng) of cDNA was amplified (first PCR) using gene-specific forward and reverse primers attached to Illumina-specific adapters for cluster generation and sequencing (5′-Illumina Nextera adaptor + sequence-specific forward and reverse primer sequence-3′, see **Table S4**). A five-cycle second PCR was performed using Phusion High-Fidelity DNA Polymerase (ThermoFisher Scientific, Waltham, MA) to incorporate Illumina Nextera primers for sample indexing. After purification with Beckman Ampure XP beads, quantification with Qubit, and quality control using the PerkinElmer LabChip GX, the indexed libraries were pooled, quantified by qPCR using the Roche KAPA Library Quantification Kit (for Illumina platforms), and sequenced on the Illumina MiSeq platform (Illumina, San Diego, CA) using paired-end 250 bp reads with a NANO flowcell and v2 chemistry reagents. Quality control of raw reads was performed using FastQC (https://www.bioinformatics. babraham.ac.uk/projects/fastqc/). Adapter trimming and quality filtering were then conducting with Trimmomatic v0.39 (78) and CutAdapt 4.1 (85) to remove Nextera adapters and primer sequences. The resulting FASTQ files were then aligned to the GRCh38 reference genome using STAR v2.7.10b (79), and alignment files were further processed with clipOverlap tool implemented into BamUtil software v1.0.15 (https://genome.sph. umich.edu/wiki/BamUtil:clipOverlap) to trim overlaps between paired reads. Pileup files and variant statistics, including coverage supporting the reference allele, coverage supporting the alternative allele, and the variant allelic fraction, were generated using PacBAM (86).

### TranSNP 3’UTR cloning

The 3’UTR regions surrounding the candidate TranSNPs were amplified from the HCT116 genomic DNA, using primers containing XbaI and EcoRI restriction sites and resulting in ∼500 bp fragments. (**Table S4**). The resulting PCR fragments were cloned immediately downstream of a modified Firefly Luciferase in pGL4.13 [Luc2/SV40] vector (Promega), where a linker sequence adding an EcoR I site (F: 5’ cAa**gaat**t**c**atatgcccgggt 3’, R: 5’ ctagacccgggcatat**gaatt** 3’) was inserted at the original Xba I site to facilitate the insertion of fragments in a desired orientation. Correct cloning was confirmed by colony PCR and Sanger sequencing.

### Plasmid DNA transfection and Luciferase assay

HCT116 cells were seeded into 96-well plates and transiently transfected the empty pGL4.13 modified plasmid or those containing the allele-specific 3’UTRs fragments along with the Renilla control luciferase (pRL-SV40) in A 3:1 ratio (60 ng plus 20 ng) using Lipofectamine^TM^ LTX and Plus^TM^ Reagent (ThermoFisher Scientific). After 48 hours, Firefly and Renilla luciferase activities were quantified using a Dual-Luciferase™ Reporter (DLR™) Assay System (Promega). The transfected cells were lysed with 50ul 1X Passive Lysis Buffer for 15 minutes. The lysates were resuspended, and 15ul of the resuspension was transferred to a white 384-well plate. Subsequently, 15ul of LAR (Luciferase assay reagent) was added, and the luminescence was read at the Varioskan LUX Microplate reader. Following this, 15ul of the Renilla substrate was added, and the Luminescence was measured again to quantify Renilla luciferase expression. RNA (TRIzol Reagent, Invitrogen) was extracted from the lysates, and the Firefly Luciferase mRNA was quantified by qPCR for additional normalization.

### CRISPR/Cas9-mediated knock-in

Edited HCT116 clones at the SNP sites were generated exploiting a CRISPR-Cas9 mediated knock-in with the Cas9 delivered as a ribonucleoparticle (RNP), as previously described (22). We designed a single gRNA (**Table S5**) in a position as close as possible to the tranSNP site to cut both alleles and maximize the efficiency of Homology-directed Repair (HDR) (**Figure S2A)**. The same protocol with few modifications (not providing the donor DNA as a template for repair and not inhibiting DNA-PKcs) was used beforehand to test the cutting efficiency of the guide RNA (gRNA). For both protocols, the gRNA was prepared by mixing 1:1 Alt-R® CRISPR-Cas9 crRNA (IDT^TM^) and Alt-R® CRISPR-Cas9 tracrRNA (IDT^TM^). The mixture was heated at 95°C for 5 minutes and let cool down at room temperature for 5 minutes for optimal annealing. 3 µl of gRNA were further incubated with 120 pmol of recombinant Cas9 for 20 minutes at room temperature to generate the ribonucleoprotein (RNP). In parallel, previously seeded HCT116 wild-type cells were harvested by trypsinization, and 2.5 x 10^5^ cells per condition were electroporated in the 4D-Nucleofector^TM^ System (Lonza) using the SE Cell Line 4D-Nucleofector^TM^ X Kit S (Lonza) as described in (22). The electroporation mixture combined 20 µl of cell suspension, 1.2 µl of Alt-R® Cas9 Electroporation Enhancer (IDT^TM^), 5 µl of RNP complex, and PBS to a final volume of 30 µl. In addition to these, 1.2 µl of Ultramer® DNA Oligonucleotide (100 µM, IDT^TM^) were added in the case of knock-in generation to provide a template donor DNA for HDR. Electroporated cells were recovered from the strip and seeded in a 12-well plate format either untreated or, in the case of knock-in generation, treated with 1 µM of NU7441 DNA-PKcs inhibitor for 48 hours to favor HDR occurrence. The media, both with and without the drug, was changed after 24 hours. The cells were kept in culture up to full recovery and expanded until 2.5 x 10^5^ cells could be harvested for DNA extraction and Sanger sequencing. DNA was extracted by adding 20 µl of the QuickExtract™ DNA Extraction Solution (Lucigen) to the cell pellet. After vortexing, the sample was incubated at 65°C for 15 minutes and 98°C for 10 minutes. 1 µl of this DNA was used for the PCR amplification using the GoTaq® Green Master Mix (Promega) in 25 µl of reaction. PCR primers (**Table S4**) were specifically designed to produce 300-700 bp long amplicons and to have the SNP of interest asymmetrically located 100-200 bp from one or the other end of the amplicon, to have optimal sequencing interpretation. The PCR product was then purified using the MinElute^TM^ PCR Purification Kit (QIAGEN). 75 ng of purified PCR products were sequenced using the Mix2Seq Kit and Sanger sequencing service provided by Eurofins. The TIDE software (Brinkman et al, 2014) was employed to identify and quantify insertions and deletions (indels) in the bulk edited population. The rs1053639 sgRNA presented a cutting efficiency of around 80%. The Synthego ICE software (Synthego Performance Analysis, ICE Analysis. 2019. v3.0) was employed to assess the knock-in efficiency in the edited bulk population. A limiting dilution assay was performed on the knock-in edited bulk population by seeding 1 cell/well in 96-well plate format to eventually obtain clones. The media was changed once a week for 3-4 weeks, then potential clones were expanded to 24-well plates, and at the same time, pellets were harvested for DNA extraction and sequencing. The efficiency of clone production was about 1 in 3 seeded cells; 85 clones were screened by sequencing.

### cDNA synthesis and qRT-PCR

RNA was extracted by using the TRIzol® reagent (Invitrogen^TM^) or the NucleoSpin ® RNA kit (Machery-Nagel) based on the downstream applications. RNA purity and quantity were assessed by NanoDrop^TM^ Spectrophotometer (Thermo Fisher Scientific), followed by random primer-mediated reverse transcription using the RevertAid First Strand cDNA Synthesis Kit (Thermo Fisher Scientific) with 1000 ng of RNA as input. The cDNA was diluted 1:8 and 12.5ng -total RNA equivalent-in 10µl reaction were used in the following RT-qPCR analysis with the qPCRBIO SyGreen Mix (PCR Biosystems). All reactions were performed in 384-well plates using the QuantStudio^TM^ 5 Real-Time PCR System (Applied Biosystems®) instrument. Primers were designed using Primer Blast (NCBI) and are listed in **Table S4**. All kits were used following the manufacturer’s instructions. All qPCR reactions were performed in triplicate, and Cq values were averaged. The fold change was calculated using *dCt* and normalized to at least two gene references.

### RNA half-life measurement

3 × 10^5^ HCT116 cells from edited clones were seeded in a 6-well plate format. After 24 hours, cells from the first timepoint (t0) well were collected by trypsinization. The cell pellet was resuspended in 500 ul of TRIzol® reagent (Invitrogen^TM^) and frozen at -80°C. The media was aspirated from the remaining wells and replaced with 10 ug/ml Actinomycin D in 2 ml of fresh media to achieve RNA polymerase II inhibition. After 20 minutes, 40 minutes, 1 hour, 2 hours and 4 hours from the treatment, cells were pelleted from the corresponding timepoint wells and resuspended in TRIzol®. Once all the pellets were recovered and frozen, they were thawed at room temperature for 15 minutes and RNA was extracted following the TRIzol® manufacturer’s instructions. RNA was quantified using the NanoDrop^TM^ Spectrophotometer (Thermo Scientific^TM^) and 1000 ng of timepoint t0 RNA from each HCT116 clone was retro-transcribed using the RevertAid First Strand cDNA Synthesis Kit (Thermo Scientific™) with random hexamers. For timepoints > t0, RNA was retro-transcribed volumetrically to the reference t0 for each clone. The cDNAs were diluted 1:8 and 2 µl in 10 µl of reaction and were used in the following RT-qPCR analysis with the qPCRBIO SyGreen Mix (PCR Biosystems) and the QuantStudio™ 5 Real-Time PCR System (Applied Biosystems®) instrument. Primers are listed in **Table S4**. GAPDH and 18S rRNA levels were also evaluated as stable mRNAs. The mRNA abundance was calculated relative to the start of transcription inhibition (t0) for each HCT116 clone (2^-dCt). The mRNA decay rate was calculated by a nonlinear regression curve fitting using the one-phase decay function (GraphPad Prism version 10).

### Western blot analysis

1.6 × 10^5^ HCT116 cells from edited clones were seeded in a 6-well plate format. When cells reached 70 - 80 % confluence, they were treated with 100 nM Thapsigargin or vehicle for 4 hours. Cells were washed with PBS, and total cellular protein was isolated using a RIPA buffer containing protease inhibitor cocktail (cOmplete™, Mini Protease Inhibitor Cocktail, Roche). Lysates prepared for analysis of phosphorylated proteins were supplemented by a phosphatase inhibitor cocktail (Phosphatase Inhibitor Cocktail I, MedChemExpress). After vortexing and incubation at 4°C for 20 min, cell lysates were centrifuged at 15000 rpm at 4°C for 15 min, with supernatant collected and stored at -80°C. Prior to western blotting, the lysates were denatured by boiling at 95°C for 10 min in 1x Laemmli Sample Buffer (Bio-Rad). The protein concentration in each sample was determined using Pierce™ BCA Protein Assay Kits (Thermo Fisher Scientific). Equal amounts of protein from each lysate were then subjected to SDS-PAGE and electrophoretically transferred to a nitrocellulose membrane (Cytiva Amersham™ Reinforced Nitrocellulose Blotting Membrane). Each membrane was blocked with 5% dry milk in phosphate-buffer saline and 0.1% Tween-20 (PBST) for 1 h and then incubated with the indicated primary antibodies overnight at 4°C. The membranes were rinsed and incubated with peroxidase-conjugated secondary antibodies for 1 hour at RT. Immunoblotted membranes were developed with ECL^TM^ Select western blot detection reagents (Amersham Biosciences, UK) and imaged on the ChemiDoc^TM^ XRS imaging system (Bio-Rad). Image processing and densitometric quantification of the bands were performed using the Image Lab software. Protein expression was normalized against housekeeping protein, which was an internal control of protein loading. Relative protein expression was then calculated by dividing the normalized intensity of the condition sample by the control. Antibodies used for western blot analysis are listed in **Table S7**. The corresponding anti-mouse and anti-rabbit secondary antibodies were used at 1:10000 dilutions.

### Protein half-life by cycloheximide chase assay

6 × 10^5^ HCT116 cells with the three genotypes were seeded in a 6-well plate format. After 24 hours, cells from the first timepoint (t0) well were collected by trypsinization. The cell pellet was resuspended in 50 µl of RIPA lysis buffer prepared *as in the Western Blot section* and supplemented with 1X protease inhibitors (Roche). In the meantime, the media was aspirated from the remaining wells and replaced with 100 ug/ml of cycloheximide (Cayman Chemical) in 2 ml of fresh media to prevent further protein synthesis. After 10 minutes, 20 minutes, 30 minutes, and 1 hour from the treatment, cells were pelleted from the wells of the corresponding timepoints and resuspended in RIPA buffer 1X as well. Once all the pellets were harvested, proteins were extracted by centrifugation at 15000 rpm for 20 minutes at 4°C, quantified by the Bicinchoninic Acid Assay (Thermo Scientific^TM^ Pierce^TM^ BCA Protein Assay Kit), and Western Blot was performed as described earlier. 35 ug of timepoint t0 protein extract from each HCT116 clone was used in Western Blot. For timepoints > t0, the protein amounts were loaded volumetrically to the reference t0, for each clone. DDIT4 and β-Tubulin antibodies were used as primary antibodies. Mouse secondary antibody was used for both the primary antibodies. β-Tubulin was found to be a stable protein and thus was used as a loading control even in this context. Upon normalization of each DDIT4 band intensity to that of the corresponding β-Tubulin, relative protein abundance over time was calculated to the start of translation inhibition (t0) for each HCT116 clone.

### Global Translation by Click-iT® metabolic labelling of proteins

Global translation was measured by using the Click-iT™ Plus OPP Alexa Fluor™ 488 Protein Synthesis Assay Kit according to the manufacturer’s instructions. The fluorescence was measured by the ImageXpress High-Content imaging system (Molecular Devices).

### Global translation by Puromycin incorporation assay

6 × 10^5^ HCT116 cells with the three genotypes were seeded in a 6-well plate format. When the cells reached 60% confluence, they were treated with vehicle or 100nM Thapsigargin for 4 or 16 hours. 1µM puromycin was added for 30 min at 37°C prior to pellet collection. Cycloheximide 100 μg/ml was used as a positive control. The cells were lysed, and the puromycin incorporation was assayed by western blot as described above. Stain-free intensity was measured as the reference.

### Cell proliferation assay

HCT116 DDIT4 edited clones were seeded at 2.5 × 10^3^ per well onto 96-well plates and incubated at 37°C in standard conditions. The cell number was followed by the high content fluorescent microscope Operetta (PerkinElmer) in digital phase contrast every 24h for 96 h.

### RNA immunoprecipitation-qPCR (RIP-qPCR)

8 × 10^6^ HCT116 cells from edited clones were seeded in 150 mm cell culture dishes (Sarstedt) and harvested after 48 hours by scraping. Cells were lysed in 800 µl of mild lysis buffer (20 mM HEPES pH 7.9, 150 mM NaCl, 1% Triton-X 100, 10% glycerol, 1 mM MgCl_2_, 1 mM EGTA, 100 U/mL RNasin (Promega), 1X protease inhibitors (Roche)). After 30 minutes of incubation on ice, the lysate was obtained by centrifugation at 15000 rpm, 4°C for 10 minutes. The protein amount was quantified by the Bicinchoninic Acid Assay (Thermo Scientific^TM^ Pierce^TM^ BCA Protein Assay Kit). Meantime, 20 µl of Protein A Dynabeads (Invitrogen^TM^) per condition were washed twice with 500 ul of IPP500 buffer (20 mM HEPES pH 7.4, 500 mM NaCl, 1.5 mM MgCl_2_, 0.5 mM DTT, 0.05% NP40) and incubated with either 0.22 µg of anti-RBMX/hnRNP G antibody (#14794, 1:40, Cell Signaling Technology®) or 0.22 µg of anti-Normal Rabbit IgG antibody (#2729, Cell Signaling Technology®) in 100 µl of IPP500 buffer for 1h 30 at room temperature in a rotating wheel. After antibody conjugation, beads were further washed once with 900 µl of IPP500 buffer and twice with 900 µl of IPP150 1x buffer (20 mM HEPES pH 7.4, 150 mM NaCl, 1.5 mM MgCl2, 0.5 mM DTT, 0.05% NP-40) and once with PBS. Nonspecific binding was blocked by incubating the beads with 300 µl of 10 mg/ml Bovine Serum Albumin (BSA) for 30 minutes at room temperature in a rotating wheel. Beads were further washed three times with 800 µl of IPP150 1X buffer and incubated with 1 mg of the protein lysate in IPP150 1X buffer and 1x protease inhibitors in a final volume of 400 µl O/N at 4°C in a rotating wheel. Before the incubation with the beads, 10% of lysate in IPP150 1X buffer was isolated as protein input for Western Blot analysis, and 1% was isolated as RNA input for qPCR analysis. After O/N incubation, the flow-through (unbound lysate) was collected for Western Blot analysis, the beads were washed four times with 800 µl of IPP150 1X buffer, and 10% of the last wash was also collected for Western Blot analysis upon elution in LB 1X (Laemmli Sample Buffer, Bio-Rad) heating at 98°C for 10 minutes. Conversely, elution both for RIP samples (beads) and RNA inputs was performed by incubation with 5 µl of >600 U/ml proteinase K solution (Qiagen) and 150 ul of Proteinase K buffer (20 mM Tris-HCl pH 7.4, 10 mM MgCl_2_, 0.2% Tween-20) at 55°C for 30 minutes in a lab heat block shaking at 700 rpm. The output RNA is extracted with TRIzol® reagent (Invitrogen) according to the manufacturer’s instructions. The total amount of eluted RNA was retrotranscribed using the RevertAid First Strand cDNA Synthesis Kit (Thermo Scientific) and 1 µl of cDNA in 10 µl of reaction was used in the following RT-qPCR analysis with the qPCRBIO SyGreen Mix (PCR Biosystems) and the QuantStudio 5 Real-Time PCR System (Applied Biosystems®) instrument. Primers are listed in Table S4.

### RNA Electromobility Shift Assay (REMSA)

The interaction between DDIT4 mRNA and RBMX protein was assessed *in vitro* by REMSA. The full-length recombinant RBMX protein was purchased from Origene (TP710403). FAM-labelled fluorescent RNA probes (GAGGGACUGAUUCC<u>U/A</u>GUGGUUGGAA-3’FAM) for the T and A alleles were purchased from Metabion. The binding reaction was set up in 10 µl in the following binding buffer (10 mM HEPES, 50 mM KCl, 1 mM EDTA, 0.05% NP-40, 10% Glycerol, 0.5 mM DTT, pH 11.5). 10 U of RNasin (1 µl) were also added to each reaction and incubated for 30 minutes at RT in the dark. Upon incubation, 1 µl of 6X Orange Dye was added to each reaction to facilitate the visualization of the samples. The binding complex was resolved by running this reaction on a 6% TBE polyacrylamide native (non-denaturing) gel at 80V for 1 hour in TBE 1X at RT. The gel was pre-run in cold TBE 0.5X at 100V for 30 minutes and imaged using the SYBR filter in the ChemiDoc^TM^ XRS imaging system (Bio-Rad).

### Cell Fractionation

Cell fractionation was performed using a 0.1% Igepal buffer in PBS 1X as in the REAP fractionation protocol, (87) coupled with an RNA extraction protocol (88) to improve the extraction of the RNA from the nuclei. Briefly, HCT116 TT and AA clones were seeded in 10-cm dishes. Once confluent, cells were detached by trypsinization, washed with PBS, and lysed in 900 µl of ice-cold 0.1% Igepal buffer in PBS 1X buffer for 2 minutes; 300 µl of this lysate were removed as whole cell lysate (WCL). The lysate was then fractionated by centrifugation at 12000 rpm for 40s, at RT. 300 µl of the supernatant were isolated as a cytoplasmic fraction (CYTO). The pellet was washed with 1 ml of 0.1% Igepal buffer and centrifuged at 12000 rpm for 40s, at RT to obtain the nuclear fraction. 50 µl out of 300 µl were isolated from each fraction for Western Blot analyses upon further lysis by RIPA buffer. The remaining 250 µl were used for TRIzol-based RNA extraction. To implement RNA extraction, the CYTO and NUCLEI fractions were resuspended in 1 ml of TRIzol. Lower volumes of TRIzol did not successfully extract RNA in our conditions. RNA was incubated for 10 minutes in TRIzol at RT, then 10 µl of EDTA 0.5 M were added before incubation at 65C for 5 min, 600 rpm (or until the pellet was completely dissolved). Subsequently, chloroform was added to follow the conventional TRIzol protocol.

### RBMX Silencing

HCT116 TT and AA cells were transfected with a single or a mixture of two out of three siRNAs targeting RBMX (IDT) at a final concentration of 40 nM for 48 hours (**Table S4**). Control cells were transfected with a non-targeting siRNA (IDT, 51-01-14-04) in the same conditions. INTERFERin (Polyplus) was used as a transfection reagent.

### Xenograft of HCT116 cells in zebrafish larvae

**Animal rearing:** Zebrafish (*Danio rerio*) strains were raised and maintained in the Model Organism Facility (MOF) at the Department CIBIO, University of Trento under standard conditions (89). The transgenic zebrafish line *Casper* (55) was used to acquire live images in transparent larvae. All zebrafish studies were performed according to European and Italian law, D.Lgs. 26/2014, authorization11/2023-PR to MM. **Larvae preparation:** Embryos were initially maintained at 28 ◦C at a maximum density of 50 embryos per Petri dish in fish water supplemented with 0.0002% methylene blue (Sigma, Burlington, MA, USA) as an antifungal agent. After 24 h, embryos were placed in fish water without methylene blue. **Xenotransplantation of human cancer cell lines:** the protocol of xenotransplantation procedure was adapted from (56,90). GFP or RFP labeled HCT116 cells (edited AA, TT, and AT clones) were harvested from a 100 mm Petri dish on the day of xenotransplantation and resuspended in 70 μl of PBS. 0.5 μL of cell suspension containing 200-400 cells was transplanted in the swim bladder of 2 dpf zebrafish larvae anesthetized in 0.16 mg/ml PBS/Tricaine (MS-222) under a Nikon SMZ800 stereomicroscope, using a micromanipulator (Marzhouser Wetzkar MM 33 links) and the manual microinjector (CellTram® 4r Air Eppendorf). Transplanted embryos were maintained at 33°C in fish water. Larvae were selected for good transplantation profile using a Leica MZ 10F fluorescence stereomicroscope. **Live-imaging and image analysis:** each larval xenograft was imaged one-day post-transplantation (D1) and 3 days post-transplantation (D3) using Leica TCS SP8 laser scanning confocal system (CD7) (Objectives: 10x/1). Images were acquired in z-stack mode with a z-step interval of 7 μm to a range of 170-230 μm depending on the tumor size. The 488/633 nm lasers were used with appropriate filters to acquire green and red fluorescent cancer cells. Image analysis was performed using Fiji software. To study tumor growth of cancer cells *in vivo*, the maximum projection of every image was created, and a threshold was applied for each larva on day 1 (D1) and day 3 (D3) to generate a mask of the tumor area. The area on D3 was normalized to the area of D1 to assess the percentage of tumor growth for each larva.

### Single-molecule inexpensive RNA fluorescence in situ hybridization

The smiFISH protocol was adapted from (91). Cells were grown on 22 × 22 mm glass coverslips (Prestige) previously stripped with 1 M HCl, sterilized, and stored in 100% ethanol. Cells were fixed in 4% paraformaldehyde (PFA) (Electron Microscopy Science) for 20 min at room temperature and then washed two times with 1× PBS. Permeabilization was performed in 0.5% Triton X-100, 1× PBS for 5 min at room temperature. Once permeabilized, samples were incubated at room temperature in a solution containing 1× SSC and 15% formamide, for 40 min and placed cell-side down on a 50-μl drop of hybridization buffer containing the probe [1× SSC, 15% formamide, bovine serum albumin (BSA) (4.5 mg/ml), 10.6% dextran sulfate, 2 mM Vanadyl Ribonucleoside Complex (VRC), yeast tRNA (0.4 mg/ml), 1 μl of probe mix] in an air-tight hybridization chamber protected from light overnight at 37°C. The probe mix preparation is described in the next paragraph. The following day, coverslips were washed twice with prewarmed 1× SSC solution containing 15% formamide for 30 min at 37°C. After that, coverslips were rinsed twice with 1× PBS, cells were stained with DAPI (1 ng/μl) (4′,6-diamidino-2-phenylindole) in 1× PBS, for 12 min at room temperature and washed with 1× PBS. Samples were mounted on precleaned glass slides using ProLong Diamond Antifade Mountant (Thermo Fisher Scientific). **smiFISH probe preparation:** probe preparation was performed as described by (91). For DDIT4 detection, 3 primary probes targeting the DDIT4 sequence were used and prepared as a 20 μM equimolar mixture in milliQ water. Sequences of the probes are shown in Table S4. Primary probes were mixed with 50 pmol of fluorescently labeled secondary probes in 1× NEB3 solution [100 mM NaCl, 50 mM tris-HCl (pH 8), 10 mM MgCl2] in a final volume of 10 μl. Primary probes were annealed with secondary probes in a PCR thermocycler (85°C for 3 min, 65°C for 3 min, 25°C for 5 min. As described in the previous paragraph, the probe mix (1 μl) was used for the smiFISH hybridization buffer preparation. **smiFISH combined with IF:** the smiFISH technique was performed first, using the protocol described in the smiFISH paragraph, and followed by the immunofluorescence protocol as described hereafter. After the overnight probe hybridization, cells were washed twice in prewarmed 1× SSC, 15% formamide solution. After two rinses in 1× PBS, cells were incubated in blocking solution (1× PBS, 2% BSA, 0.1% Triton X-100) for 3 hours and incubated for 1 hour at room temperature with the primary antibody diluted in blocking solution. For RBMX detection, we used the antibody in Table S7 at 1:200 dilution. Cells were then washed five times for 6 min with 1× PBS, 0.1% Triton X-100, and incubated with the appropriate fluorescent secondary antibody for 1 hour [goat anti-rabbit Alexa Fluor 488 (AF488), from Thermo Fisher Scientific, catalog no. A-11034, 1:1500 dilution; donkey anti-mouse AF647, from Thermo Fisher Scientific, catalog no. A-31571, 1:1000 dilution]. After incubation, cells were washed five times for 6 min with 1× PBS, 0.1% Triton X-100, and stained with DAPI (1 ng/μl) in 1× PBS for 12 min. After two washes with 1× PBS, coverslips were mounted on glass slides using ProLong Diamond Antifade Reagent (Thermo Fisher Scientific). **smiFISH/IF image acquisition by confocal microscopy:** images were acquired using a confocal laser scanning Leica TCS SP8 inverted microscope (Leica Camera AG) with a plan apochromatic 63×/1.40 oil immersion objective and 2× zoom. Normal photomultipliers (PMTs) were used to detect DDIT4 smiFISH signal, RBMX, b-tubulin, and DAPI signals. Optimal acquisition parameters, including nanometer range of excitation, gain, offset, and laser power, were optimized for the visualization of each fluorophore at the beginning of the acquisition and maintained throughout each experiment. Z stack images were acquired in each experiment with a 0.3-μm step and 2048 × 2048 pixel size resolution using Leica Application Suite X (LAS X) imaging software.

### Confocal image analysis

Images were analyzed using CellProfiler 4.0.7 (Broad Institute, Inc.). Briefly, cell nuclei were identified as the primary object using the DAPI signal. For each object, the cellular region was defined by propagating from the nucleus, applying a low segmentation threshold to the smoothed smiFISH signal. The cytoplasm was defined as a tertiary object by subtracting the nuclear masks from the cellular regions. DDIT4 smiFISH signal spots were segmented as objects with a typical diameter range of 10 to 30 pixels. Subsequently, objects were localized based on their position relative to the nuclear or cytoplasmic regions. Since the majority of the spots exhibited a perinuclear localization, the nuclear regions were reduced by 5% to prevent misleading classification. Morphological features and intensity values for both DDIT4 smiFISH and RBMX signals were calculated at the object level.

### Immunohistochemistry

Briefly, three-micron sections from paraffin blocks by seven patients with colon adenocarcinoma were cut and incubated (37°C overnight). IHC staining was performed using Bond III IHC autostainer for full Automated Immunohistochemistry (Leica Biosystems). Antigens were unmasked with Bond Epitope Retrieval Solution 1 Leica AR9961 and incubated with the following antibody (diluted with Bond Primary Antibody Diluent, AR9352 Leica): DDIT4/REDD1 at dilution 1:400 (Proteintech 10638-1-AP). Samples were stained with BOND IHC Polymer Detection Kit (DS9800) and counterstained using Hematoxylin solution (Leica). The IHC score was obtained by analyzing the slides by a certified pathologist (G.B.) and applying also ImageJ, a digital image processing software. Informed consent was obtained from all patients for the use of tissue materials for research purposes.

#### Statistical analysis

Results were plotted and statistically analyzed using GraphPad Prism, version 10. The statistical test applied, the number of replicates, and the level of significance are provided in each figure legend.

## Supporting information

Suppplementary Tables S1-S8

## Declarations

### Ethics approval and consent to participate

Informed consent was obtained from all patients for the use of tissue materials for research purposes.

### Availability of data and materials

The RNA-seq dataset re-analyzed for this study is deposited in GEO under the GSE95024 accession number.

### Authors’ contributions

MHH and LA performed most of the experiments, prepared figures and contributed to the first draft of the manuscript; TV, SV, DD, performed the RNA-sequencing analysis and developed the pipeline to identify tranSNPs; FL, performed with MHH and LA the experiments in Zebrafish and prepared the first draft of the relative methods and results; AM, VV supported the DDIT4 gene editing approaches; GPG performed with LA the smiFISH experiments; AM supported the design and execution of the cell sorting experiments; PG, MP supported all HTS-based experiments; CV, VdS, RB designed the targeted resequencing and performed RNA-sequencing experiments; VM supervised MHH in planning and performing translation assays; EC contributed to the design and the evaluation of the results of smiFISH experiments; SF, GB performed, analyzed and interpreted the IHC data; SZ supervised LA in designing and performing in vitro RBMX-RNA binding and GLORI at DDIT4 3’UTR; MM supervised and contributed to the design and interpretation of the experiments in zebrafish; LLF supervised the editing approaches, contributed to the design, visualization and interpretation of the co-culture experiments, contributed to drafting and revising the manuscript; AR supervised the study, in particular RNA-seq data analyses, preparation of Tables, drafting of the manuscript; AI supervised the study, secured the main findings, drafted the manuscript.

## Acknowledgments

We thank Dr. Ainan Geng and Prof. Hashim M. Al-Hashimi, Department of Biochemistry and Molecular Biophysics, Columbia University, NYC, USA, for the DDIT4 3’UTR RNA structural predictions. We thank Dr. Mounira Chalabi, Ribosome, Translation and Cancer Team Universite’ Claude Bernard Lyon, France, for technical support to M.H.H. We thank Alice Casata, Irene Cagol, Massimo Andreis, Micol Bottamedi, Federica Sontacchi, and Christian Cestaro for technical support during their Bachelor’s internships. We thank Dr. Ezequiel Mariano Rivero and Prof. Fatima Gebauer, Gene Regulation, Stem Cells, and Cancer Program, Centre for Genomic Regulation (CRG), Barcelona, Spain, for support with the RIP protocol. We thank Dr. Gabriella Viero, Institute of Biophysics, CNR Unit at Trento, Italy, for support with sucrose gradient fractionation and data analysis.

## Supplementary Figures and Tables

**Figure S1.**
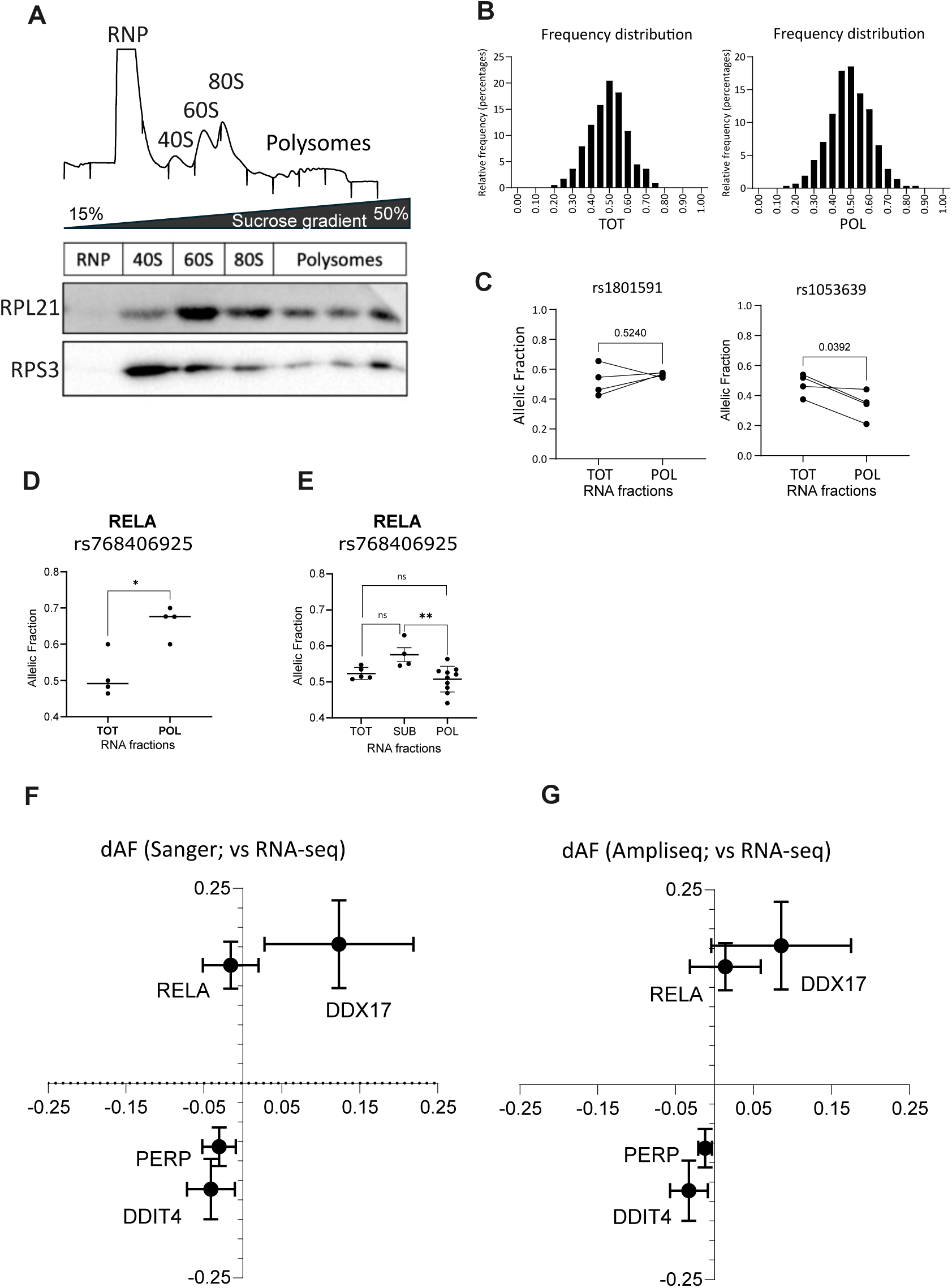
TranSNPs identification and validation -related to Figure 1-. **A)** Representative polysome profile of HCT116 cells and western blot results for markers of the small (RPS3) and large (RPL21) ribosomal subunits used as controls for sedimentation. Lysates separated to equilibrium density by ultracentrifugation on a linear 15-50% sucrose gradient were fractionated following UV absorbance recorded at 254 nm. The fractions representing polysomes (POL) that were pooled together for subsequent analysis are indicated. **B**) Distribution plotted as the percentage of the relative frequency of average allelic fractions for all the heterozygous SNPs that could be studied in the RNA-seq experiment, for total (left figure) and polysomal (right figure) RNA. AF data for each of the four experimental replicates is provided in Table S1. The RNA-seq dataset is deposited in GEO (accession number: GSE95024). **B)** Examples of paired t-test statistics exploited to assess the concordance of differential AF measures across pairs of polysomal and total RNA biological replicates. **D**) Plots of the allelic fraction (the number of alternative allele reads divided by the sum of alternative and reference allele reads) measured for rs768406925 in the RELA gene from RNA-seq results obtained in total cytoplasmic lysates (TOT) or pooled polysomal fractions (POL). **E**) Allelic fraction measurements obtained by Sanger sequencing and electropherogram quantification from independent biological replicates of polysome profiling experiments in HCT116 cells for rs768406925 in RELA. **F**) Correlation between delta allelic fractions measured for the SNPs in the four indicated genes using RNA-seq (Y-axis) or Sanger electropherograms (X-axis). The average and standard deviations of replicates are plotted. **G**) Correlation graph as in F, focusing on the RNA-seq data (Y-axis) and the results obtained by Ampliseq-based targeted resequencing (X-axis).

**Figure S2.**
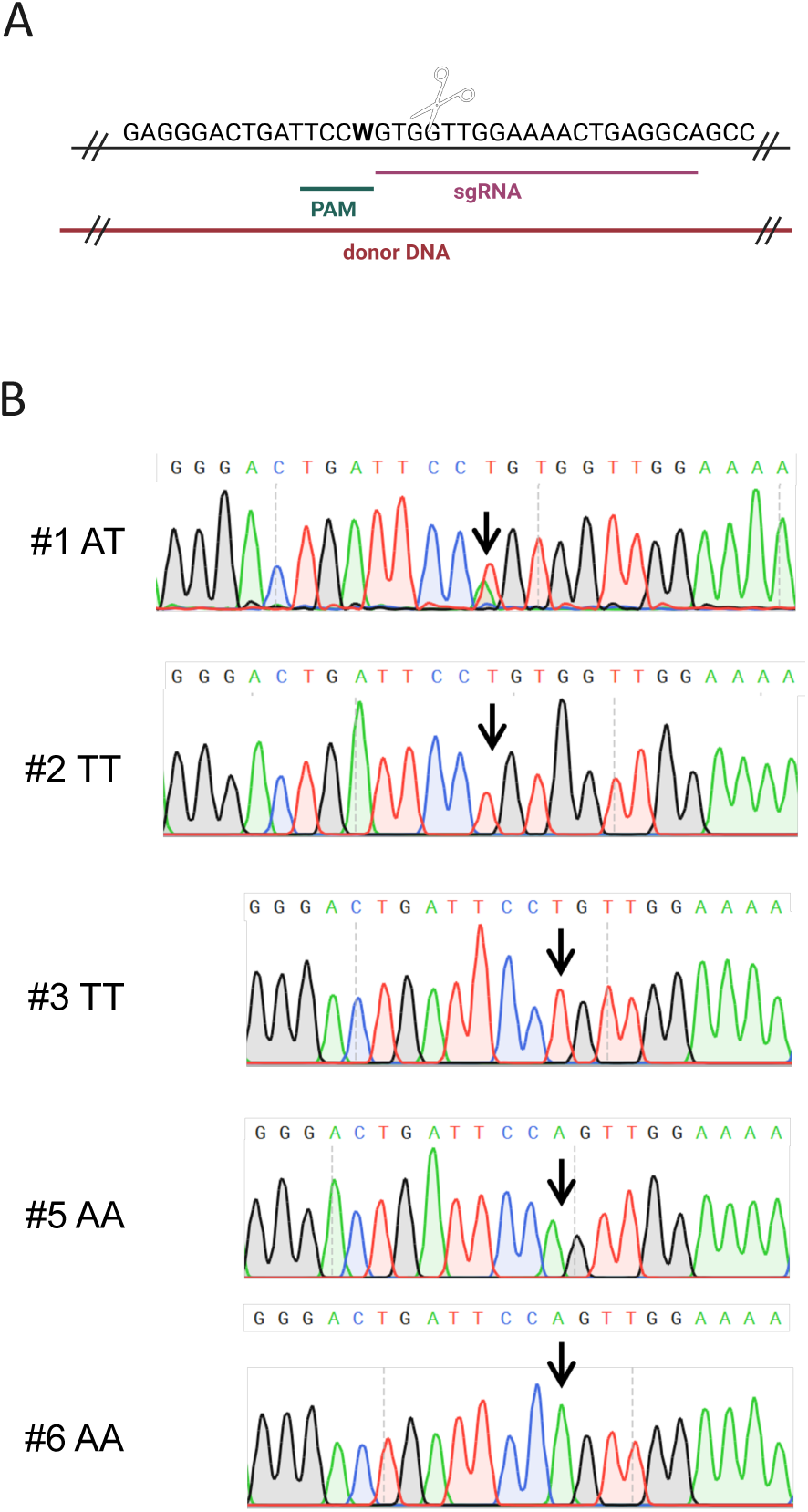
Editing of HCT116 cells to obtain the three genotypes for the rs1053639 T/A SNP. Parental HCT116 cells - heterozygous for the rs1053639 SNP - were edited to obtain homozygous clones for the A and the T alleles. **A)** CRISPR/Cas9-mediated knock-in was employed for this aim. We annealed a gene-specific crRNA with a tracrRNA, obtaining a sgRNA for DDIT4. The same sgRNA cut both alleles since the SNP represented the “N” base of the -NGG PAM. An 85-nt long donor DNA with the endogenous sequence (mixture of 50% A and 50% T) was purchased to be used as a template. **B)** Representative electropherograms of Sanger sequencing analyses indicating the genotype of the selected clones for rs1053639.

**Figure S3.**
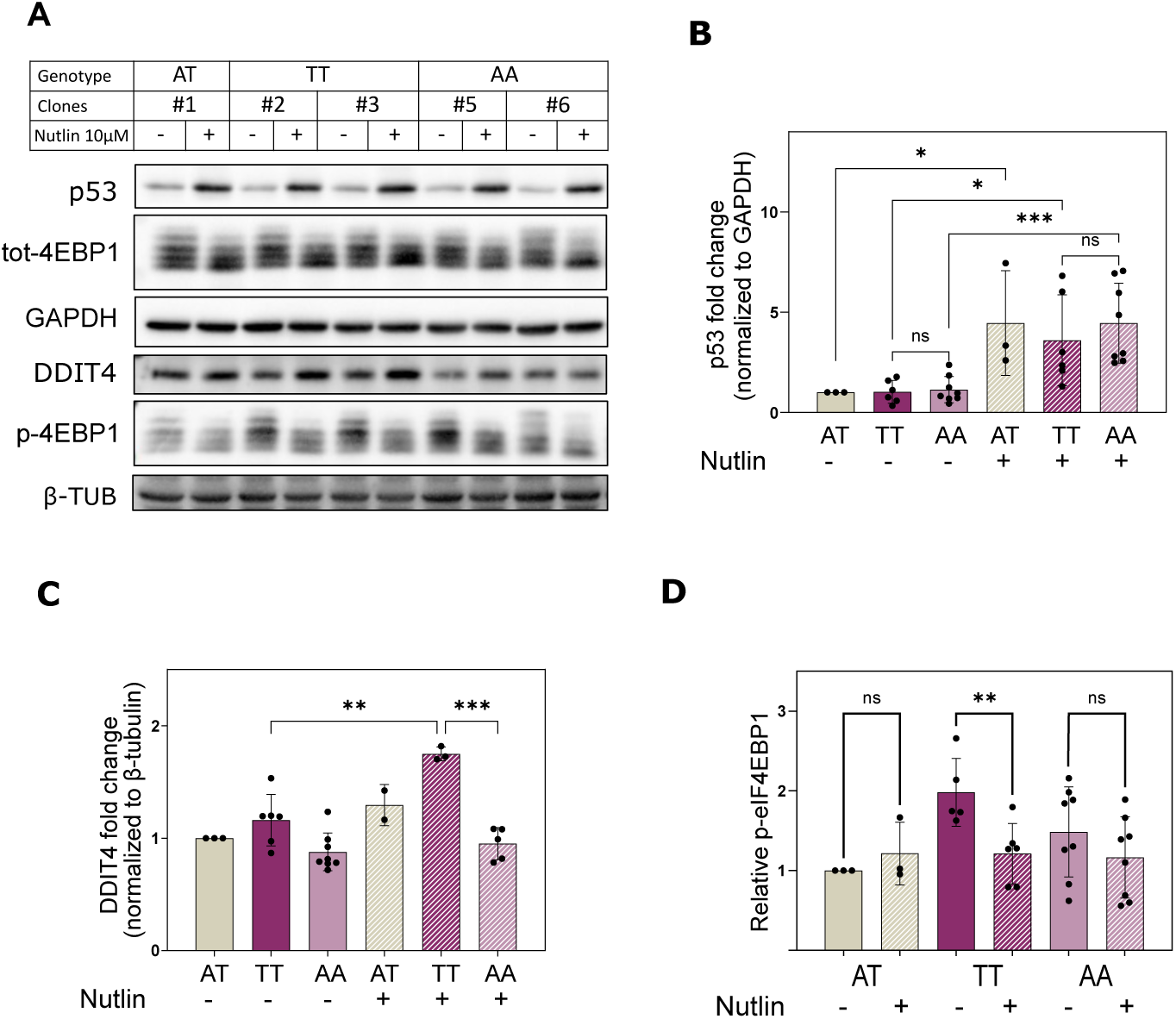
The rs1053639 tranSNP genotype impacts on mTOR pathway markers upon Nutlin. treatment. HCT116 parental cells and clones differing for the rs1053639 homozygous genotypes were treated with either Nutlin 10 µM or vehicle control for 16 hours. **A)** Cell lysates were analyzed by immunoblotting for the induction of p53 and DDIT4 protein levels and the phosphorylation state and levels of the eIF4EBP1 mTORC1 target. β-Tubulin and GAPDH were used as references. **B, C)** Densitometry analysis of p53 and DDIT4 protein levels shows significant induction upon Nutlin treatment. **D)** Densitometry analysis of phosphorylated eIF4EBP1 blots reveals that significant inhibition of the phosphorylation of eIF4EBP1 is apparent only in the rs1053639 TT homozygous clones. ***p-value <0.005. **p-value <0.001. *p-value <0.05. ns, not significant. unpaired t-test.

**Figure S4.**
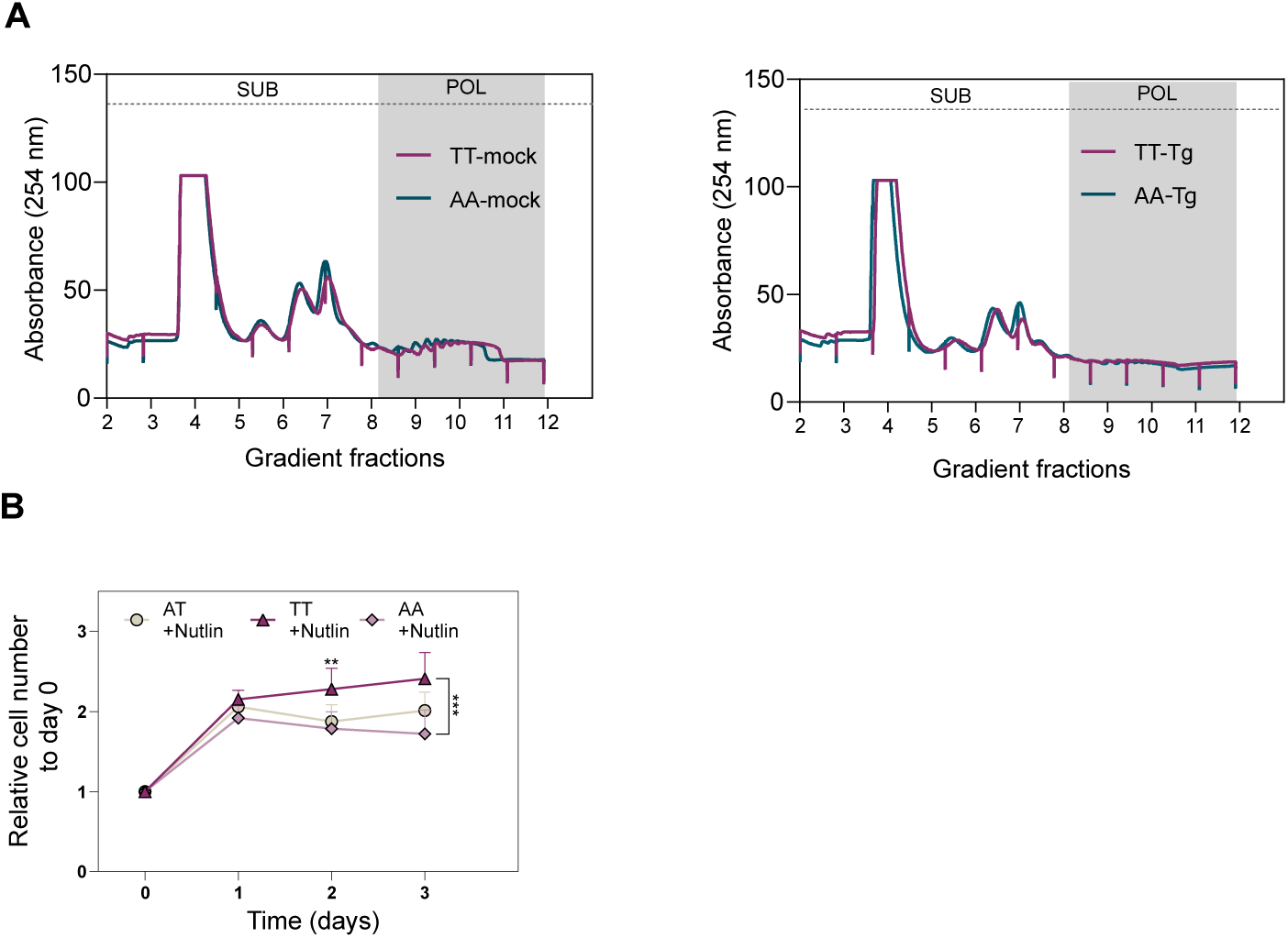
The DDIT4 rs1053639 genotype does not strikingly impact global translation and cell proliferation in response to cellular stresses. **A)** Representative polysome profiles of HCT116 TT and AA clones treated with vehicle (Left) or thapsigargin 100nM for four hours (Right). Lysates separated by ultracentrifugation on a linear 15-50% sucrose gradient were fractionated following UV absorbance recorded at 254 nm. The fractions representing RNA complexes used for subsequent analysis are indicated in the figure. **B)** HCT116 cells with the three genotypes differing for the rs1053639 were treated with 5µM Nutlin for 72 hours. The sensitivity of the clones was followed every 24 hours by the Operetta HCS system. Relative cell number curves were plotted as the mean of several replicates ± standard deviation (SD). ***p-value <0.0005. Ordinary two-way ANOVA.

**Figure S5.**
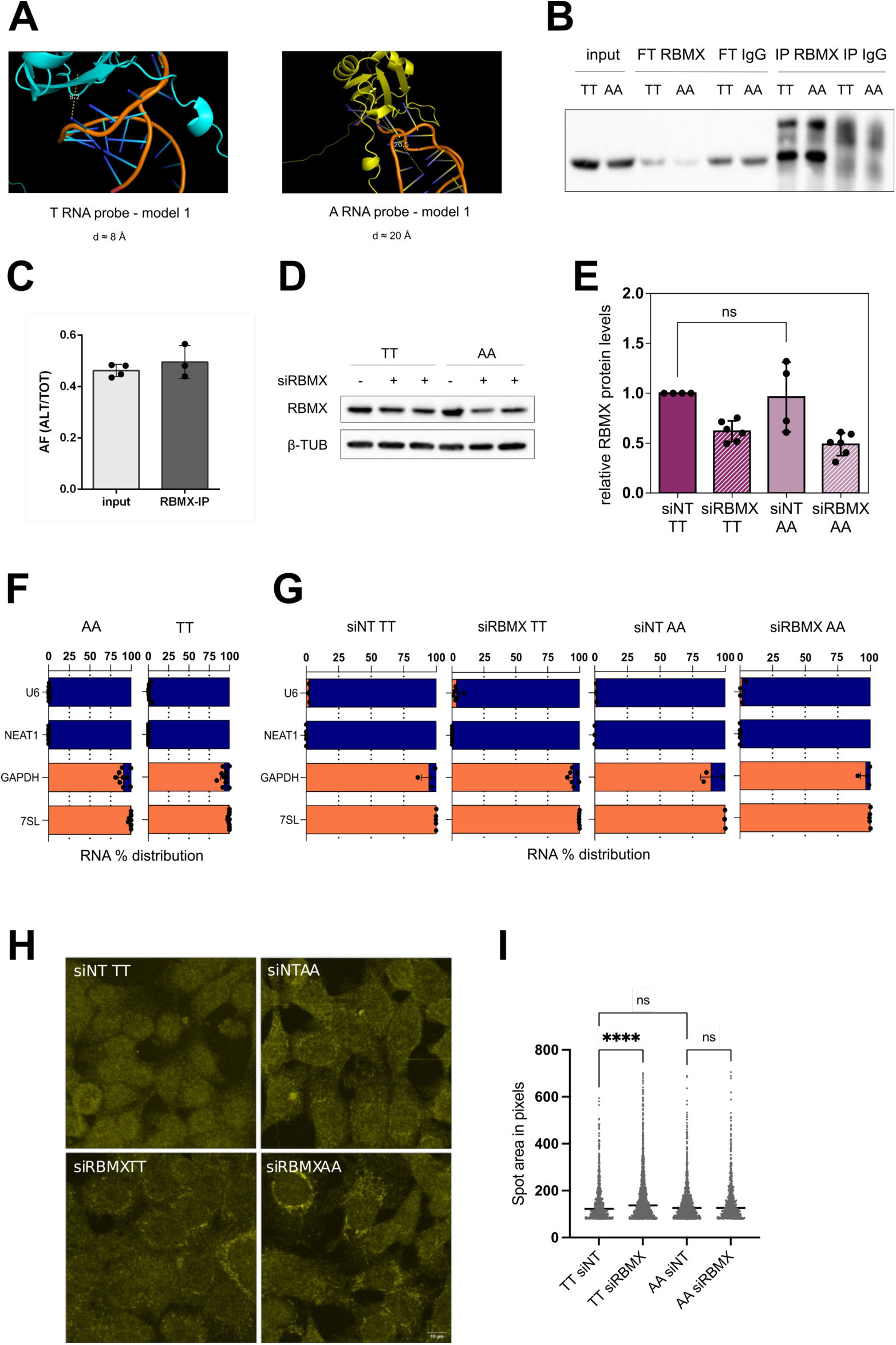
The rs1053639 tranSNP genotype impacts mRNA fate through the RBMX protein. **A)** AlphaFold3 was used to predict the interactions between the full-length RBMX protein and the RNA probes (A, T) used for the RNA EMSA. Four models were explored; model 1 is shown as representative. The tool allows the calculation of the distance between a selected RNA nucleotide and a selected protein portion. The distance between the T SNP nucleotide and the RBMX protein was lower than the distance between the A SNP nucleotide and the same protein in all four models. **B)** RBMX was specifically immunoprecipitated across TT and AA cells with similarly high efficiencies. An IgG control antibody of the same species as the anti-RBMX antibody was used as a control for specificity. FT: flow-through, unbound lysate upon IP. **C)** The endogenous RBMX protein was immunoprecipitated either from a whole cell lysate or from a nuclear extract in heterozygous parental HCT116 cells. RNA from the input and IP samples was extracted and reverse-transcribed. The cDNA was amplified by performing RT-qPCR and submitted to Sanger sequencing. Results are indicated as the AF of the alternative allele (A) over the Total (A+T) and represent the mean ± SD of three independent experiments. **D, E)** TT and AA clones were treated with either a mixture of siRNAs targeting RBMX or the siRNA control for 48 hours. Representative immunoblot and the densitometric analysis show that the RBMX protein is equally expressed independently of the rs1053639 and that the efficiency of RBMX silencing was similar across TT and AA clones. β-Tubulin was used as a loading control. **F)** U6 snRNA and NEAT1 were used as markers of the nuclear fraction, 7SL and GAPDH were used as markers of the cytoplasmic fraction. However, GAPDH mRNA is not completely cytoplasmic, which is consistent with previous reports. Markers for the AA cells are shown on the *left*, and markers for the TT cells are shown on the *right*. **G)** Markers for RNA fractionation upon siNT or siRBMX. U6 snRNA and NEAT1 were used as markers of the nuclear fraction, and 7SL and GAPDH were used as markers of the cytoplasmic fraction. **H)** Confocal images related to Figure 4G: smiFISH of a TT and an AA cell clone after control (siNT) or RBMX depletion, showing only the DDIT4 mRNA, in yellow. Scale bar is 10 μm. **I)** Area of perinuclear, cytoplasmic DDIT4 mRNA spots in pixels. **** p<0.0001, ordinary one-way ANOVA. ns = not significant.

**Figure S6.**
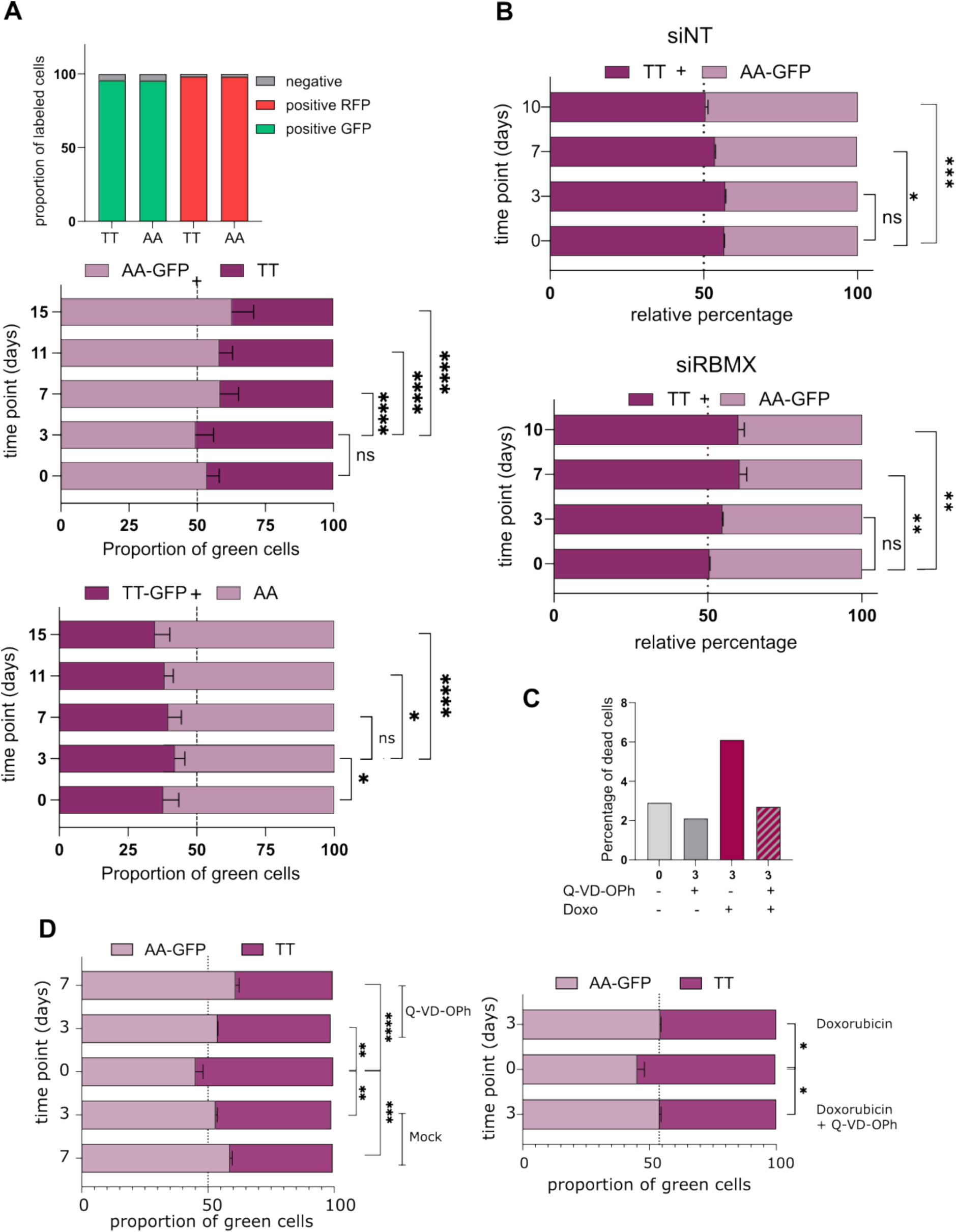
The rs1053639 tranSNP AA genotype shows a competitive growth advantage in prolonged culture conditions. **A)** (Top) DDIT4 clones harboring the TT and AA genotypes were both separately transduced with Lentiviral particles to constitutively express either EGFP or MiRFP. After transduction, cells were treated with 5 µg/ml blasticidin for at least 15 days, to reach at least 95% of GFP+ or RFP+ cell populations. (Middle and Bottom) TT and AA clones were co-cultured using both combinations for the fluorophore. The graphs plot the relative proportion of cells and are representative of at least four replicates for each combination. ns = not significant; *p-value < 0.05; ****p-value <0.0001, ordinary two-way ANOVA. **B**) Relative percentage of TT and AA cells at time zero, when the co-culture was started, and at the indicated time points. The relative proportions were calculated by detaching the cells and counting 10,000 events by FACS, identifying GFP-positive (AA) and GFP-negative cells. At day 0, day 3, and day 7, cells were transfected with a control siRNA (siNT) (Top) or an siRNA targeting RBMX (Bottom). The average relative percentage and the standard deviations of three replicates are shown. The statistical analysis over day 0 is shown. ns = not significant; *p-value <0.05; **p-value <0.001; ***p-value <0.0001, ordinary two-way ANOVA. **C)** HCT116 clones were treated with Doxorubicin (3µM) and/or Q-VD-OPh (10µM) and tested after three days. Percentage of dead cells measured by FACS in the indicated conditions. **D**) The two graphs represent the proportion of GFP+ cells (AA) at day 0 and at the indicated time points of co-culture and treatments. *p-value <0.05; **p-value <0.001; ***p-value <0.0001; ****p-value <0.0001, ordinary two-way ANOVA.

**Figure S7.**
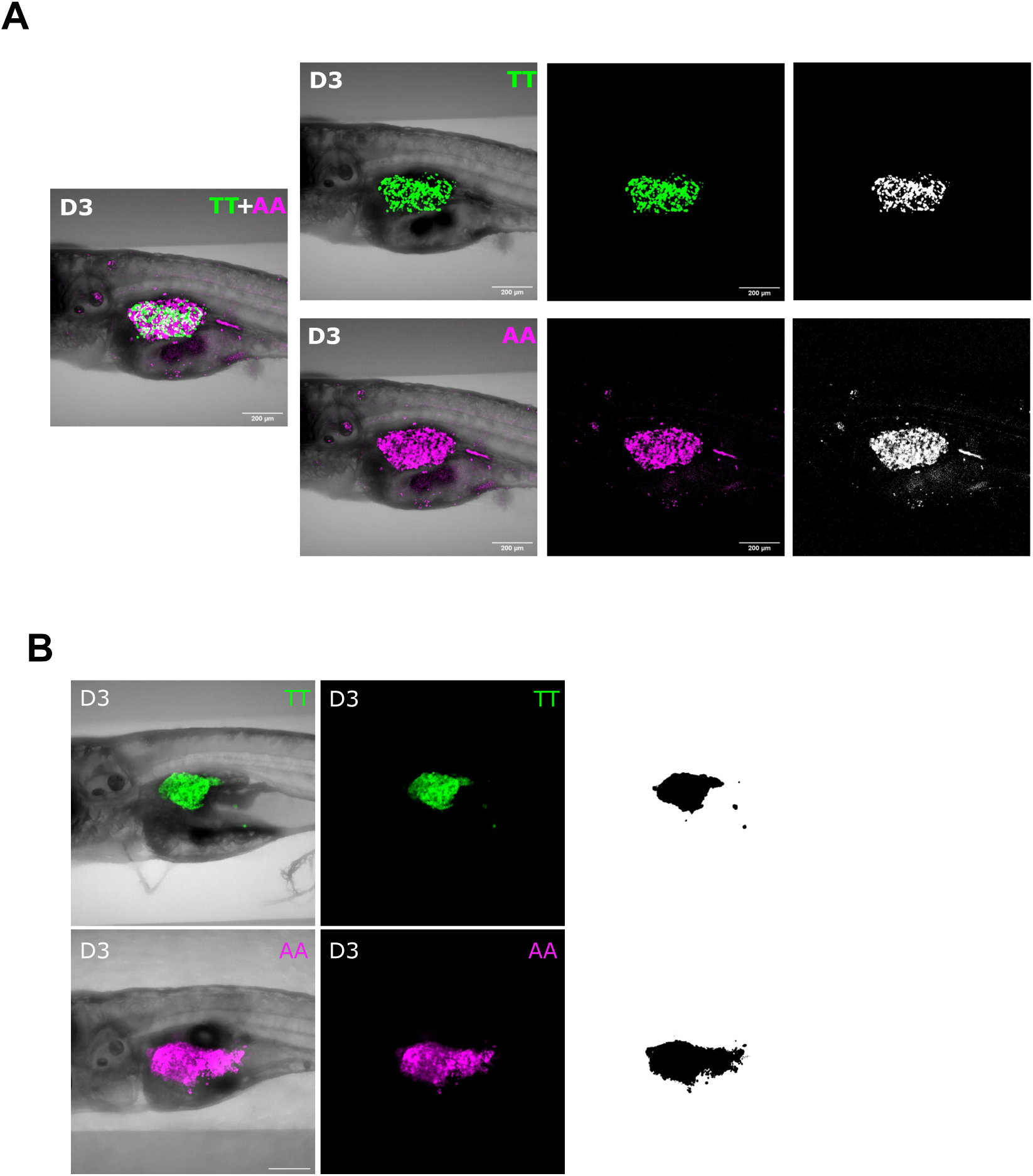
DDIT4 rs1053639 impacts tumor growth and genotypic competition in zebrafish larvae xenotransplants. Image analysis was performed by Fiji software, and quantification of the tumor growth was carried out by applying fixed thresholds for fluorescent intensity, as described in Materials and Methods. **A, B)** Representative images for both confocal and its corresponding Fiji quantification mask.

**Figure S8.**
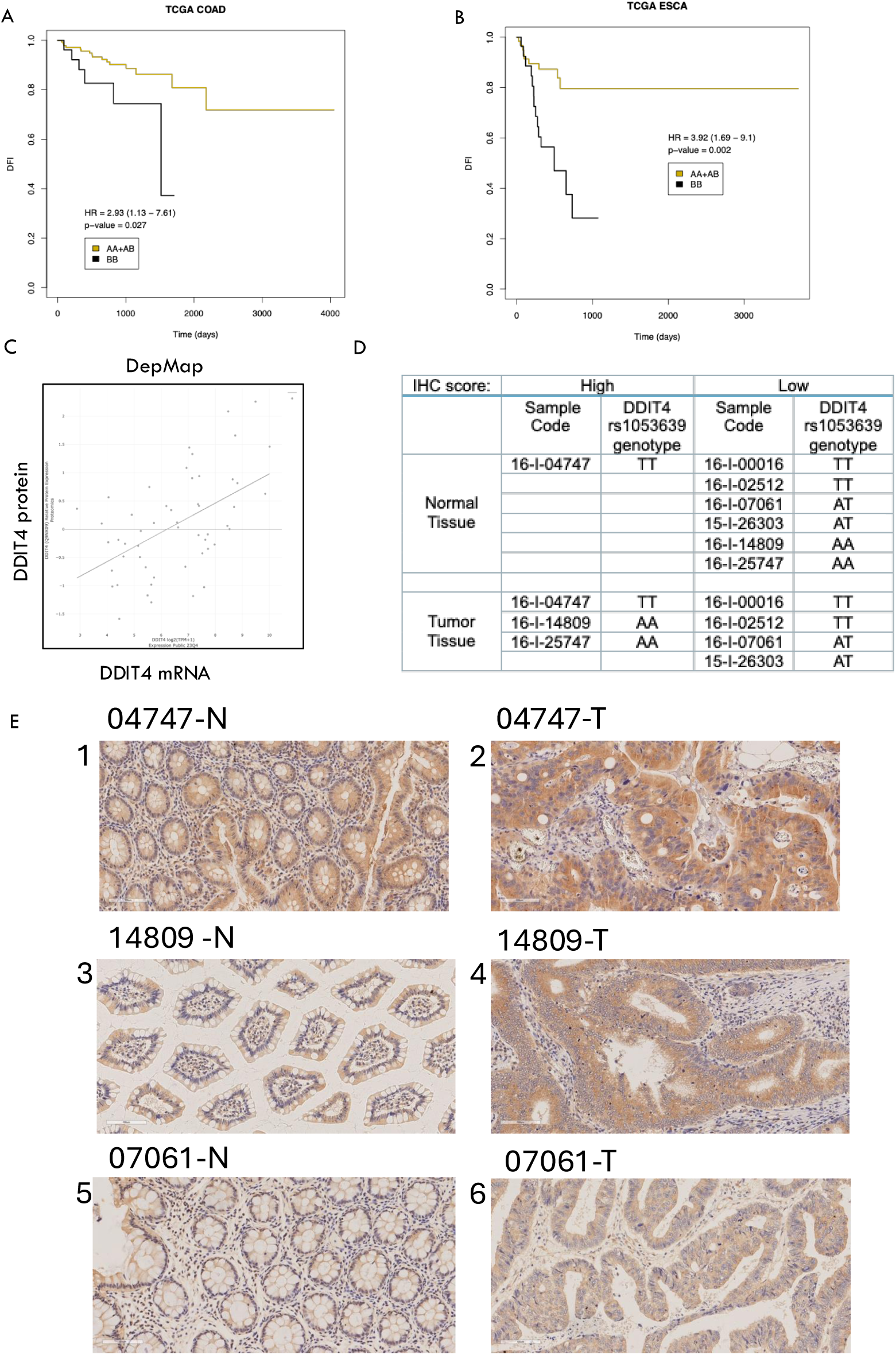
Exploration of the prognostic value of rs1053639 genotype. **A-B)** Survival probability from TCGA data plotted after stratifying patients based on the rs1053639 genotype and under a recessive model (A = reference allele (T); B = alternative allele (A)). **C**) Correlation between DDIT4 protein and mRNA levels in cancer cell lines from the DepMap database. **D)** Genotyping results for rs1053639 in seven colon cancer samples. The IHC score was obtained by analyzing the slides by two certified pathologists and applying a custom script in Fiji -see methods-. **E)** Representative images of IHC DDIT4 staining. -N and -T after the sample code refer to normal colon mucosa or cancer tissue, respectively. The sensitivity of IHC staining was verified by paraffin-embedding pellets from our edited HCT116 cell clones (not shown). Images of sample 04747-N and -T, genotype TT, show high expression of cytoplasmic DDIT4 (E1 and E2). Sample 14809-N, genotype AA, shows a low expression of cytoplasmic DDIT4 (E3), while the 14809-T tumor counterpart shows high expression (E4). Finally, images of sample 07061-N and -T, genotype AT, both show low cytoplasmic expression of the DDIT4 protein (E5 and E6).

## List of Supplementary Tables

**Table S1**. List of analyzable heterozygous SNPs in HCT116 RNA-seq dataset

**Table S2**. List of the tranSNPs identified. ID, gene name, location, and AF data are reported

**Table S3**. RNA-binding protein binding predictions for rs1053639.

**Table S4**. Sequences of primers and siRNAs

**Table S5**. Sequence of crRNA and donor DNA sequences and summary of the efficiency of the CRISPR/Cas9-mediated HDR.

**Table S6**. Summary table of the genotypes of selected rs1053639 edited clones.

**Table S7**. Table of the antibodies used in this study.

**Table S8**. Comparison of the expression of tranSNP host genes in colon adenocarcinomas and controls, based on TCGA data.

